# Temporally discordant chromatin accessibility and DNA demethylation define short and long-term enhancer regulation during cell fate specification

**DOI:** 10.1101/2024.08.27.609789

**Authors:** Lindsey N. Guerin, Timothy J. Scott, Jacqueline A. Yap, Annelie Johansson, Fabio Puddu, Tom Charlesworth, Yilin Yang, Alan J. Simmons, Ken S. Lau, Rebecca A. Ihrie, Emily Hodges

## Abstract

Epigenetic mechanisms govern the transcriptional activity of lineage-specifying enhancers; but recent work challenges the dogma that joint chromatin accessibility and DNA demethylation are prerequisites for transcription. To understand this paradox, we established a highly-resolved timeline of DNA demethylation, chromatin accessibility, and transcription factor occupancy during neural progenitor cell differentiation. We show thousands of enhancers undergo rapid, transient accessibility changes associated with distinct periods of transcription factor expression. However, most DNA methylation changes are unidirectional and delayed relative to chromatin dynamics, creating transiently discordant epigenetic states. Genome-wide detection of 5-hydroxymethylcytosine further revealed active demethylation begins ahead of chromatin and transcription factor activity, while enhancer hypomethylation persists long after these activities have dissipated. We demonstrate that these timepoint specific methylation states predict past, present and future chromatin accessibility using machine learning models. Thus, chromatin and DNA methylation collaborate on different timescales to mediate short and long-term enhancer regulation during cell fate specification.

## INTRODUCTION

Normal cell differentiation depends on the coordinated regulation of lineage-specifying gene enhancers to drive transcriptional programs. Epigenetic mechanisms mediate this process on multiple levels, from DNA methylation (DNAme) to chromatin accessibility (ChrAcc). Canonical models of gene regulation assume that both ChrAcc and DNA demethylation are inherent to gene transcription. However, we and others have demonstrated that DNAme and chromatin dynamics are not as tightly linked as previously thought, challenging the causal relationship between DNAme, gene enhancer regulation and transcription.^1–3^

DNAme has been classically defined as transcriptionally repressive, playing an essential role in transposable element silencing and heterochromatin formation.^4–9^ Whole genome methylation data across distinct cell types and developmental stages have shown that, whereas most of the genome is methylated, hypomethylated regions denote gene regulatory elements.^10–16^ Promoters are largely hypomethylated across cell types, while hypomethylation of enhancers is cell-type specific and differentiation-dependent.^17–20^ Accordingly, gene enhancers commonly acquire both ChrAcc and DNA hypomethylation to promote transcription of lineage-specifying genes; but whether these two epigenetic changes occur on similar timescales or how the timing of demethylation affects enhancer function relative to accessibility is unknown.

Previous studies report that TET oxidase activity, rather than passive demethylation, is responsible for establishing hypomethylation at most enhancers^21, 22^, and the by-product of TET activity, 5-hydroxymethylcytosine (5-hmC), is enriched at enhancers in embryonic stem cells.^23^ Constitutive disruption of TET activity results in cell differentiation defects in both embryonic and adult cells.^24–30^ For example, loss of TET2 leads to increased methylation of neural progenitor cell (NPC) enhancers, delaying the induction of NPC differentiation genes.^31, 32^ Likewise, TET2 plays a specific role in hematopoiesis^33^, and loss of TET2 leads to transcriptional skewing of hematopoietic stem cells.^34^ DNAme restricts the binding of certain transcription factors (TFs) to DNA;^29, 35–41^ thus, failure to demethylate lineage-specifying enhancers precludes the expression of critical genes, blocking cell differentiation cascades.^42^

Despite these important findings, prior work comparing steady state data revealed transcriptionally “discordant” gene enhancers that are at once accessible and methylated or inaccessible and hypomethylated.^1, 19, 43^ Contrary to dogma, the implications of these studies are that ChrAcc and DNAme dynamics are not always concurrent and DNAme does not invariably repress enhancer activity. Moreover, in time course studies, we previously discovered that ChrAcc and gene activation occur irrespective of enhancer demethylation, and demethylation is not required for successful terminal differentiation of human macrophages.^1, 44^ Similarly, a separate study showed that gene activation precedes DNA demethylation during infection of post-mitotic dendritic cells.^45^ Whether this decoupling of DNAme, ChrAcc, and transcriptional dynamics extends to replicating cells must be determined.

The maintenance and modification of DNAme patterns are subject to the kinetics of enzyme activity and DNA replication.^22, 46^ TET initiated 5-hmC represents an intermediate state that is eventually resolved through active base-excision repair mechanisms involving thymine DNA glycosylase (TDG) or by passive dilution during replication.^47, 48^ The demethylation mechanism depends on the developmental setting. In certain cell types , replication is required for the majority of methylation loss through either passive dilution of 5-mC or its oxidized intermediates.^48, 49^ Other cell types, such as post-mitotic neurons, rely on active removal of oxidized 5-mC products entirely.^50, 51^ Moreover, demethylation mechanisms may be fully dispensable in late differentiation settings.^25, 49, 52, 53^ Thus, the observation of DNAme dynamics is likely affected by the temporal properties and mechanism of demethylation acting in the model system.

Additionally, while ChrAcc is dictated by TF binding activities, some, but not all, TF interactions with DNA are methylation sensitive.^2, 38^ In fact, some TFs bind methylated, and even inaccessible, DNA.^53^ Single molecule studies probing DNAme and TF occupancy found that only a small subset of enhancers depends on DNA demethylation for transcriptional activity.^2^ Further, dynamic transcriptional responses have been observed without DNA demethylation of regulatory sequences, suggesting transcriptional activity, at least in the short-term, supersedes DNA demethylation mechanisms.^45, 54, 55^

These collective findings highlight a contradictory understanding of how DNAme relates to ChrAcc and transcription that is, to some extent, at odds with phenotypes observed in DNAme modifier mutants. Moreover, the temporal resolution to understand the significance of mixed, and in some cases “discordant”, epigenetic states is lacking in most datasets – especially for fate-specifying enhancers experiencing epigenetic transitions. The role of DNAme on gene regulation may be time and context dependent; thus, a key to understanding the *causal* relationship between DNAme, gene regulation, and cell differentiation is to determine the timing and order of DNAme changes compared to TF occupancy, ChrAcc, and transcription.

Here, we simultaneously quantified DNAme, ChrAcc, and TF footprints from single DNA fragment libraries^56^ to construct a high-resolution timeline of their dynamics during NPC differentiation. Overall, we show a majority of lineage-specifying enhancers undergo periods of DNA demethylation that are temporally distinct from chromatin. In fact, a substantial subset of enhancers loses DNAme despite transient opening and closing of chromatin. The greatest loss in DNAme occurs several days after initial ChrAcc and transcriptional changes, primarily between two and 6 days of differentiation. Furthermore, hypomethylation of these enhancer regions persists after these activities subside. Measuring site-specific 5-hmC^57^, we identified regions and periods of active demethylation that initiate before, and continue after, TF binding, suggesting the arc of DNA demethylation from beginning to end occurs outside of TF activity. Finally, using machine learning, we show that 5-hmC accumulation forecasts future ChrAcc, while 5-mC logs past activity. Our findings clarify how enhancers are regulated on different timescales by ChrAcc and DNAme, arguing that DNAme is not a gatekeeper of transcription, but serves to stabilize enhancer transitions during cell fate specification. Understanding the timescale over which DNAme exerts its regulatory function is fundamental to interpreting the functional consequences of epigenetic patterns in normal and disease states.

## RESULTS

### Directed differentiation of HESCs to NPCs displays extensive DNA demethylation within chromatin accessibility loci

We used a well-established dual-SMAD inhibition protocol to differentiate human embryonic stem cells (HESCs) to neural progenitor cells (NPCs) (**Figure 1A**).^58^ With this system, two SMAD inhibitors, Noggin and SB431542, are applied to HESCs grown in a monolayer on Matrigel, allowing for robust, feeder-free generation of NPCs in less than two weeks. In contrast to our previous work^1^, this differentiation system has several important characteristics: 1) a longer differentiation timeline allows for frequent sampling of timepoints, 2) cells continue to proliferate throughout a 12-day time course, 3) NPCs retain the potential to be further differentiated into functionally specialized neural cells, and 4) the resulting cells can be characterized at each stage of differentiation using known HESC and NPC markers including Oct4, Sox1/2, Nestin and Pax6 (**Figure S1A**). Finally, using single cell RNA-seq for a subset of timepoints (0-, 2-, and 6-days post-induction), we observed cell clustering by timepoint. Within each time point, no distinct subclusters were observed, indicating homogeneous/synchronous differentiation of cells and ruling out cell heterogeneity as a potential confounder in our results, especially for genomic regions with mixed epigenetic states (**Figure 1B**).

**Figure 1:**
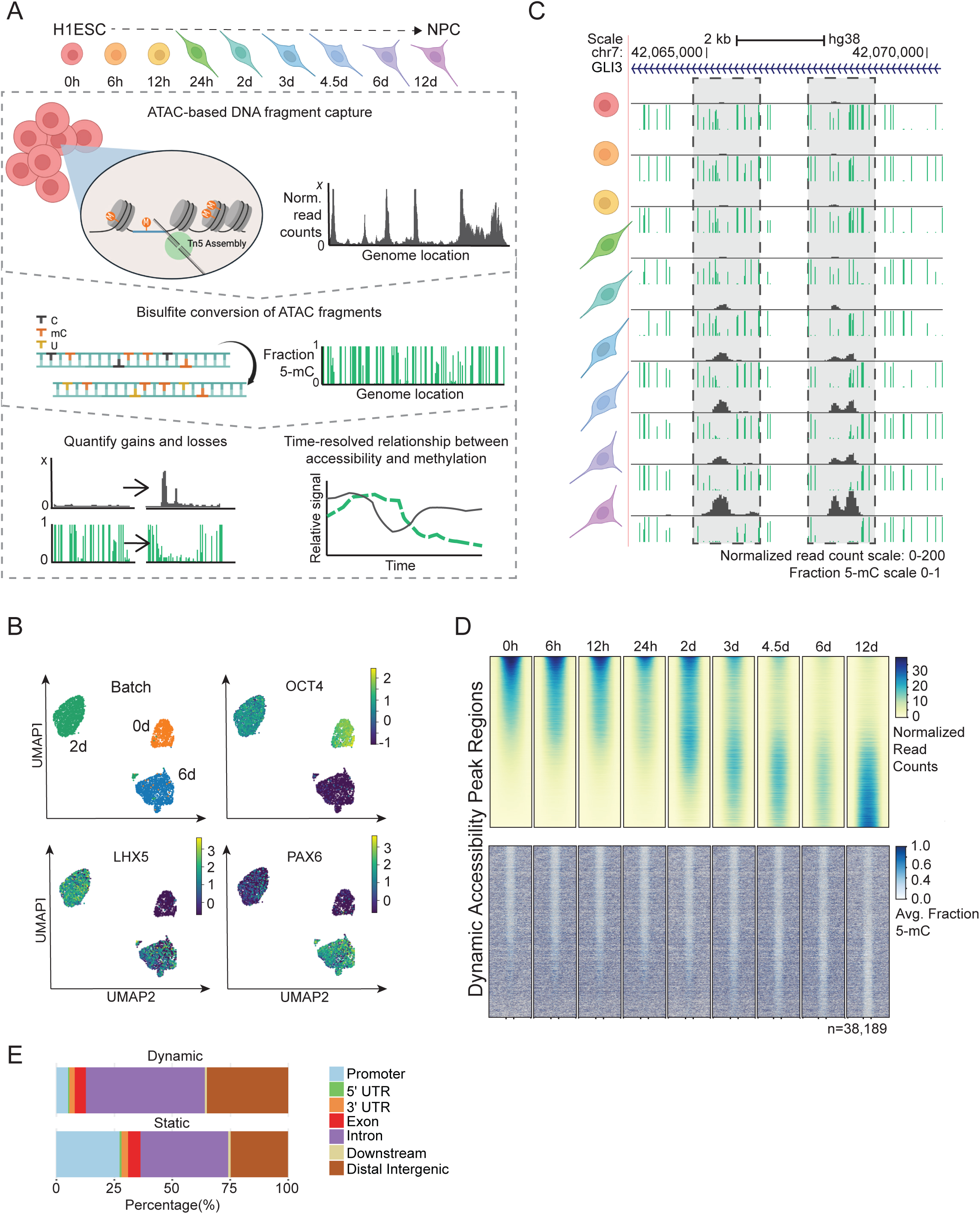
Directed differentiation of HESCs to NPCs displays extensive DNA demethylation within chromatin accessibility loci. (A) The experimental design of ATAC-Me consists of four main steps. HESCs are differentiated to NPCs for 12 days and samples are taken at nine time points throughout the differentiation process. DNA fragments are isolated from Tn5 accessible chromatin followed by sodium bisulfite conversion to quantify methylation state of open chromatin regions. Analysis of resulting data captures dynamic behaviors of DNAme and ChrAcc over time. (B) UMAPs of single cell RNA-seq data for samples analyzed at 0, 2 and 6 days of differentiation. Groups (Batches) segregate according to timepoint and homogeneously express markers of ESCs (OCT4), intermediate NPCs (LHX5), and differentiated NPCs (PAX6). Marker gene overlays are scaled by normalized and transformed read count values. (C) UCSC Genome Browser tracks display ATAC-Me derived DNAme and ChrAcc measurements at the GLI3 locus. Grey boxes highlight two regions that gain accessibility and lose DNAme. The fraction methylated reads at each CpG site is represented by the height of the green bar. Accessibility is represented by normalized read counts shown in grey. Both tracks are merged signal of two replicates. (D) Heatmaps display the ChrAcc and DNAme signal of all dynamic ChrAcc peaks at each time point. Regions are sorted by decreasing normalized read count signal intensity at the 0-hour time point. Regions are scaled to 500 bp and plotted along the center of each +/- 0.5 kilobases and 1 kilobases for ChrAcc and DNAme, respectively. (E) Proportion of dynamic (n=38,189) and static (n=63026) regions annotated to genomic region classes is shown. Related to Figure S1.

We performed ATAC-Me and bulk RNA-seq in parallel for two biological replicates of nine timepoints following NPC induction, including 0 hours, 6 hours, 12 hours, 24 hours, 48 hours, 3 days, 4.5 days, 6 days, and 12 days (**Figure 1A, Table S1-2**). These timepoints were chosen to capture early, intermediate, and late events in the gene regulatory cascade as well as transient ChrAcc and DNAme states. For all timepoints, ATAC-Me and RNA-seq replicate libraries were reproducible and showed similar sequence complexities (**Figure S1B-D**; Spearman ρ: 0.86-0.98).

Capturing ChrAcc and DNAme from a single DNA fragment source with ATAC-Me combined with deep sampling of timepoints permits quantification of their relationship with high spatiotemporal precision (**Figure 1C**). Initial genome-wide analysis identified a total of 101,215 chromatin accessibility loci from all time points collected. The majority of these loci remained static and open for the duration of the time course (n=63,026), whereas a substantial subset (n=38,189) displayed dynamic accessibility over time (**Figure 1D, S1E**). Dynamic regions are predominantly located in intronic and intergenic genomic locations (∼85%) where cell specific gene enhancers typically reside, while static regions locate to a greater degree near promoters, where accessibility is stable across cell and tissue types (**Figure 1E**).

Contrary to data obtained from terminally differentiated (and post-mitotic) hematopoietic cells^1^, we captured extensive DNAme changes within these dynamic ChrAcc regions (**Figure 1D, S1E**). This result is expected given the differentially methylated regions previously identified from comparisons of steady state HESCs and NPC methylomes^16^, as well as the length of the time course and the extent of reprogramming required to achieve the cell phenotype transition in this model system. However, our initial analysis further demonstrates that, whereas chromatin accessibility changes are bidirectional, DNAme changes are not. Many early hypomethylated regions remain hypomethylated despite closing chromatin, and most opening sites lose rather than gain DNAme. Altogether, our approach reveals new insights regarding the unique timing of these epigenetic transitions, the direction of change, and the regulatory elements involved at a scale and resolution that have not been previously determined.

### Unsupervised clustering of chromatin accessibility reveals temporally distinct regulatory groups with divergent changes in enhancer states

To identify temporal patterns across individual chromatin accessibility loci, we performed unsupervised clustering on the 38,189 dynamic regions using normalized read counts for each time point (**Figure 2A**).^59^ Using a combination of methods to determine the optimal number of C-means groups (**Figure S2A-C**), we defined seven clusters each containing unique accessibility regions that track closely with the nine selected time points (n=3929-7520 regions). Within 6 hours after differentiation induction, there are notable changes in chromatin accessibility and each subsequent timepoint is associated with a specific cluster of accessibility regions, illustrating how rapidly and transiently chromatin responds to differentiation signals.

**Figure 2:**
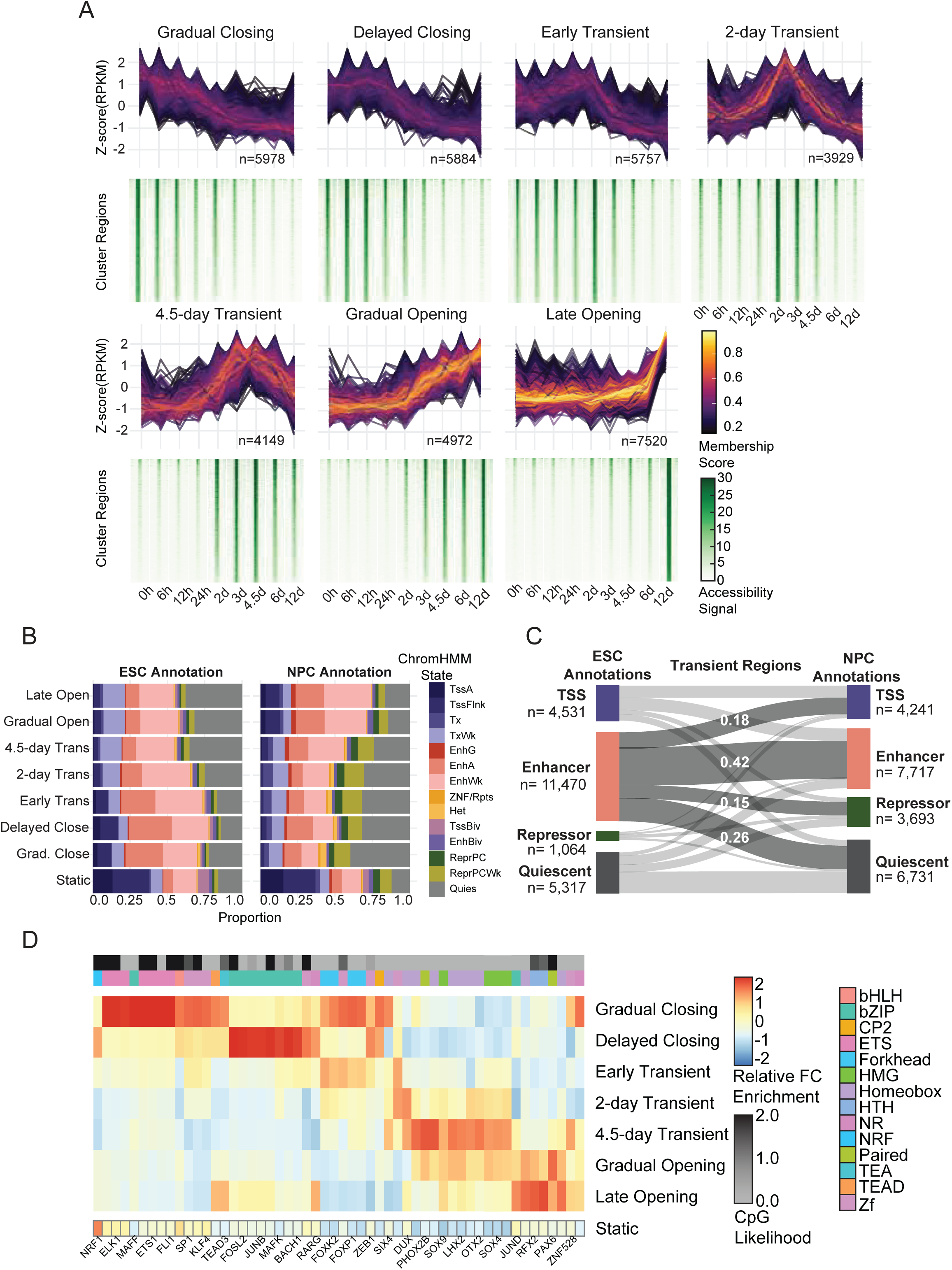
Unsupervised clustering of chromatin accessibility reveals temporally distinct regulatory groups with divergent changes in enhancer states. (A) ChrAcc regions with differential accessibility over time (|log_2_-fold| > 2, adjusted p-value < 0.05) were clustered using fuzzy C-means clustering. The standard difference of normalized ATAC-Me signal intensity (z-score) over time for each region within a cluster is shown, with line color representing the membership score defined by that cluster. Heatmaps displaying the normalized accessibility signal across the cluster regions for each timepoint are shown below. Heatmaps are sorted by decreasing normalized read count signal intensity at the 0-hour time point for each cluster. The region count for each cluster is displayed. (B) Chromatin state annotations of cluster regions using the chromHMM^61^ 18-state annotations from HESCs and NPCs. The proportion of regions in each state for the cluster is displayed for all dynamic and static regions. (C) A Sankey plot displays the change in regions’ chromatin states from the ESC to NPC stages for all *Transient* regions. (D) Motif enrichment was performed for each dynamic ChrAcc group using HOMER. The relative enrichment (z-score of enrichment values across all dynamic clusters) of the topmost variable TFs are shown and are filtered for motif redundancy. For a comprehensive list, see Table S3. The enrichment score of the same motifs in static regions is also shown. TF family is displayed as an annotation column along with CpG content likelihood. CpG likelihood in each TF consensus motif is calculated as described in Motto^97^. Related to Figure S2.

Chromatin accessibility represents one of the first steps in the regulatory cascade of enhancer regulation^60^, and we show that chromatin accessibility occurs in multiple waves over the time course; thus, we classified each cluster into three major categories: *Opening*, *Closing*, and *Transient*. These broad classifications can be further separated by specific temporal behaviors. The *Gradual Closing* cluster contains approximately 6,000 regions which begin closing almost immediately while *Delayed Closing* regions remain open for the first 12-24 hours (**Figure 2A**). The *Transient* groups each reach peak accessibility at different times but close by 12 days. *Gradual Opening* and *Late Opening* regions are both open at the NPC stage, but the rate of accessibility differs with *Gradual Opening* regions undergoing a gradual increase where *Late Opening* regions do not become accessible until 6 days post induction.

The temporal resolution of our time course enables dissection of accessibility dynamics and assignment of gene regulatory elements to discrete stages of HESC-to-NPC differentiation. Accordingly, each dynamic accessibility cluster is enriched for gene ontologies that draw a clear distinction between early, transient, and late events such as negative regulation of developmental processes like circulatory system development (early), neuron differentiation (transient), and forebrain and cerebral cortex development (late, **Figure S2B-C**). By contrast, static regions are enriched for genes involved in general housekeeping processes (**Figure S2C**).

Overlap of dynamic regions with 18-state chromHMM annotations^61^ trained on data from either HESCs or NPCs revealed substantial overlap of enhancer and repressor states with dynamic regions compared to static regions (**Figure 2B**). Comparing the ESC chromHMM to NPC chromHMM annotations for the same regions shows that, in *Opening* regions, enhancer annotations increase substantially while quiescent annotations are lost (**Figure S2D**). *Transient* and *Closing* regions undergo substantial switching from enhancer states in HESCs to repressor and quiescent states in NPCs (41% and 45%, **Figure 2C, S2E**).

Motif enrichment analysis revealed strong correspondence between distinct sets of TF motifs and time-point associated accessibility clusters (**Figure 2D, Table S3**). These TFs include canonical pluripotency factors like Oct4/Sox2/Nanog in *Closing* regions and NPC marker Pax6 in *Late Opening* regions. *Transient* regions demonstrated staggered opening and closing dynamics, suggesting short-lived TF activity within those regulatory elements. The *4.5-day Transient* regions, for example, are enriched for Otx2, a TF shown to drive neural fate during early differentiation.^62^ In total, we observed 14 different TF families that defined the sequence content of the cluster behaviors.

Given that CpGs are the major substrate for DNA methylation, we considered the CpG content of each accessibility cluster. Whereas static regions have a higher CpG density (observed/expected∼0.4, **Figure S2F**) supported by their higher CpG island promoter content, dynamic regions display a range of CpG densities (mean obs/exp=0.174-0.50, **Figure S2F**). We determined whether CpG density could be attributed to specific TF motifs, finding that CpG containing TF motifs were associated with *Opening* and *Closing* clusters, rather than *Transient* regions (**Figure 2D**). This apparent dearth of CpG containing motifs in *Transient* clusters is supported by the significantly lower CpG density in these regions compared to *Opening* (p-value <2e-16) and *Closing* (p-value=3.06e-9) clusters, suggesting an underlying link between sequence and methylation kinetics (**Figure 2D, S2G**).

### DNAme dynamics are unidirectional and temporally discordant with chromatin accessibility

To gain a detailed understanding of the temporal relationship between ChrAcc and DNAme, we quantified DNAme of regions within each accessibility group for every timepoint (**Figure 3A**). These data revealed that, whereas chromatin and transcriptional changes begin as early as 6 hours post-induction, notable changes to DNAme do not begin until 48 hours (**Figure 3A, S3A**). Overall, open regions that remain constant are constitutively hypomethylated throughout the time course (*Static* regions, **Figure 3A**). Among the dynamic accessibility regions, many display “concordant” changes with DNAme, where decreases in DNAme accompany increases in ChrAcc (**Figure 3B-C, S3B**). In fact, DNAme loss is the most prevalent pattern across all dynamic regions; however, unlike rapid and transient changes in ChrAcc that occur in both directions, the greatest loss of DNAme occurs during a distinct window of time between 2-6 days (**Figure S3A-B**). This delay creates a subset of regions that pass through a “discordant” state in which they are open and methylated during enhancer activation.

**Figure 3:**
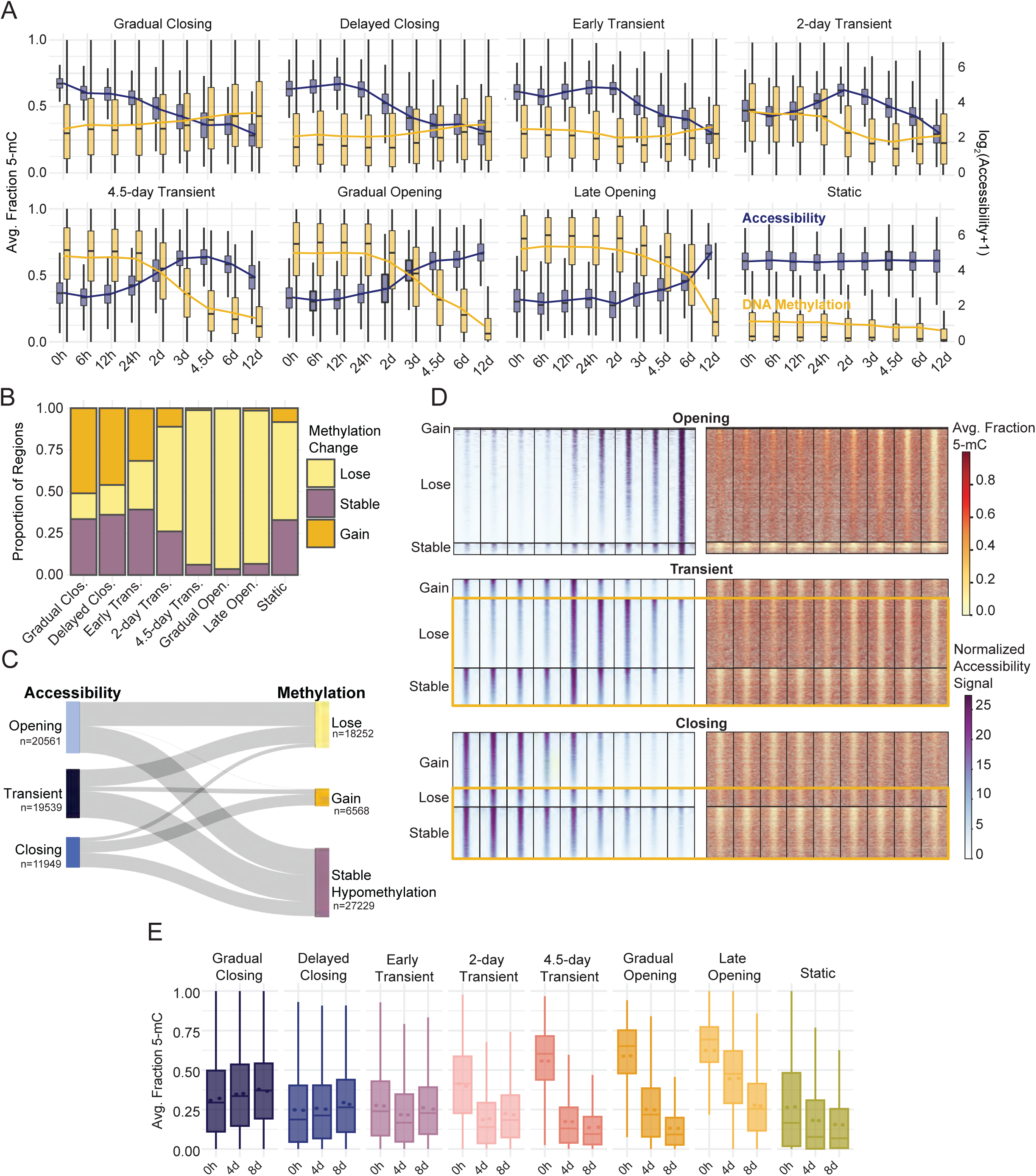
DNAme dynamics are unidirectional and temporally discordant with chromatin accessibility. (A) Dual-axis boxplots of accessibility signal distribution (normalized read counts, blue) for each timepoint grouped by dynamic TCseq clusters. A pseudocount is added and the displayed data is log transformed for display. The corresponding average fraction methylation distribution across each region group and timepoint is shown in gold. The boxplots display the median of the signal distribution, and the line overlay represents the average signal at each timepoint. (B) The proportion of regions within each accessibility cluster that experience a gain, loss or no change in methylation over time. Regions were grouped based on the change of average regional methylation values over the entire time course, 0 to 12 days. The stable methylation group represents those regions which showed a change less than 10% between the 0 hour and 12-day time point. Methylation classification of “lose” or “gain” indicates a change of at least 10% in the average methylation between the 0 hour and 12-day timepoints in either direction. (C) The temporal relationship between accessibility and methylation behaviors represented by a Sankey plot. Accessibility subgroups represent dynamic regions from all TCseq clusters. Clusters were grouped by their dominant accessibility trend (i.e., opening, transient and closing) while the methylation classification from (B) was maintained. (D) Regional methylation and accessibility are displayed for all dynamic accessible regions. Heatmaps are grouped by accessibility subgroup then methylation behavior, the methylation classification from (B) was maintained. Yellow boxes highlight regions which display discordant epigenetic states by the end of the time course. (E) Average fraction DNAme values determined by whole genome 6-base sequencing across regions contained in each ChrAcc cluster are shown. 6-base sequencing was performed on samples collected at 0, 4, and 8 days of differentiation. Regional methylation values represent the average fraction methylation from two biological replicates. Related to Figure S3.

Gain of DNAme was a less common occurrence in our dataset (15.3% of dynamic ChrAcc regions, **Figure 3B-C**). We hypothesized that this may be due to the slower kinetics of DNAme gain and loss. However, extended time does not result in substantial gain of methylation for newly closed regions, as demonstrated by *Closing* ChrAcc groups that remained hypomethylated after 12 days of differentiation (**Figure 3D, S3B-C**). Moreover, both *Transient* and *Closing* regions continue to lose DNAme even after the regions return to a closed state. These dynamics create another “discordant” epigenetic state whereby regions are inaccessible and hypomethylated or where regions are demethylated and remain hypomethylated despite opening and closing of chromatin (**Figure 3C-D**, **S3C**). We performed unsupervised clustering analysis on DNAme of all accessible regions to obtain groups based on similarity of their methylation dynamics rather than ChrAcc dynamics (**Figure S3D**). These data confirm that the DNAme patterns emerge independently of ChrAcc, but largely recapitulate the patterns observed when regions are clustered by accessibility.

To determine whether this observation is due to a sampling bias (DNA fragments derived from closing regions are less abundant in ATAC-Me), we performed whole genome methylation profiling using 6-base sequencing^57^, an orthogonal method to bisulfite-based sequencing, at 0, 4 and 8 days of differentiation. This approach showed high correlation with methylation measured by ATAC-Me and recapitulated the methylation patterns observed across the 7 accessibility behaviors (Pearson=0.83-0.9, **Figure 3D, S3E**).

In line with previous studies^1^, gene expression changes tracked more closely with ChrAcc than DNAme (**Figure S3F**). For many genes associated with closing clusters, expression decreased in tandem. Likewise, gene expression increased for genes proximal to opening regions. In fact, these changes occurred long before associated DNAme changes appeared. These findings suggest a general decoupling of DNAme from the ChrAcc and gene expression changes that drive the ESC to NPC transition. Overall, we observed three major types of DNAme trends during differentiation: slow response relative to ChrAcc, limited restoration of DNAme to closed enhancer regions, and continued demethylation of *Transient* and *Closing* accessible regions. The combination of these DNAme characteristics with rapid ChrAcc responses produces enhancer regions with discordant epigenetic signatures, contradicting the textbook model that DNAme (or lack thereof) is immediately synonymous with chromatin and gene expression changes (**Figure 3E**). These data also demonstrate the role of DNA hypomethylation as a record of current and historically active enhancers.

### Enhancer demethylation appears prior to, and is maintained independently of, TF binding

Using Tn5 cut site frequencies generated in the ATAC-Me libraries, we performed TF footprinting to estimate TF occupancy of dynamic accessibility regions (**Figure 4A**).^63^ We then calculated the average methylation at these binding sites for all timepoints (**Figure 4B**). We considered identified sequence motifs in the JASPAR CORE Vertebrates collection, which allowed us to reduce redundancy and consolidate patterns generated from TFs with high degrees of similarity– especially those within the same family.^64, 65^ From our timepoint-paired RNA-seq data, we determined that patterns of TF expression specifically produce analogous groups to those produced by accessibility (**Figure 4C, Table S4**). Example footprint profiles of the POU family displayed in Figure 4A include footprints of OCT4, POU3F1, and BRN2, which are expressed at different times during differentiation (**Figure S4A**). These expression profiles follow a clear switch in binding events between 2-6 days across the different accessibility regions. This switch coincides with a window during which the highest level of DNAme loss occurs and is representative of a larger trend we observe across TFs (**Figure S3**). Thus, to better predict the footprint source, we used TF expression to narrow the scope of TFs considered in our analysis. Integrating TF footprints and TF expression enabled us to calculate methylation of regions before, during, and after a predicted binding event, giving a clearer picture of the timing of regulatory changes.

**Figure 4:**
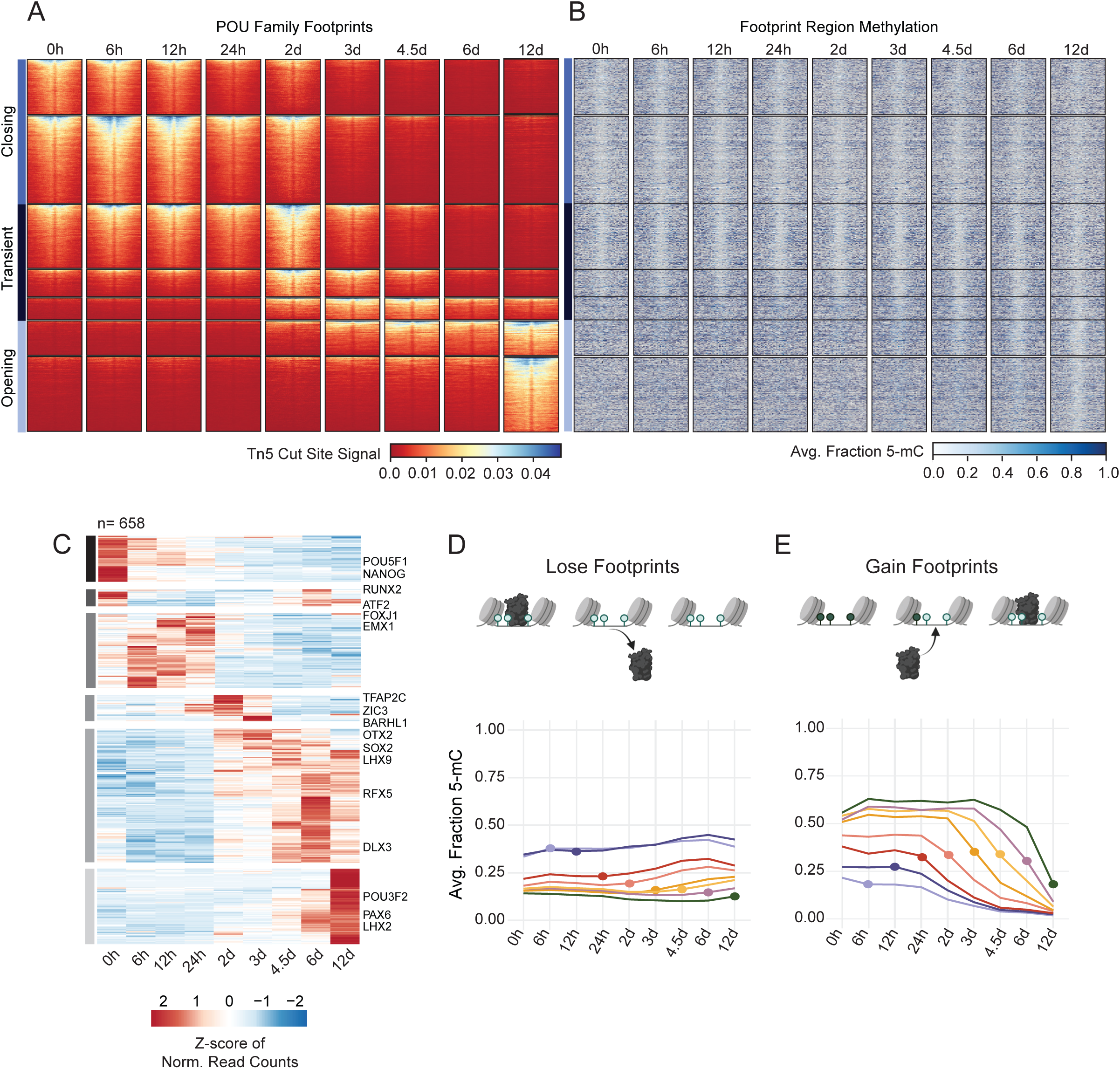
Enhancer demethylation appears prior to, and is maintained independently of, TF binding. (A) Heatmaps display cut site signal centered around TF footprint sites containing POU family motifs +/-200bp. Footprint sites are defined by POU motif sequences +/- 50bp. Regions are grouped by previously defined accessibility clusters and organized within each cluster according to descending cut site signal intensity. Horizontal bars indicate the larger subgroups defined by accessibility behavior over the time course. (B) The methylation heatmap displays the corresponding proportion methylation at each CpG site within in the footprint site with a flanking distance of +/-1kb. Regions are sorted according to (A). (C) Heatmap displays TF expression determined by RNA-seq for all TFs expressed at any time point. Normalized read counts (FPKM) are scaled by row and ordered by hierarchical clustering. Horizontal grey bars define six groups with specific temporal expression patterns. Select TFs are labeled to the right of their respective rows. (D, E) Line plots show average regional methylation values over time visualized by TF binding behavior. The dot represents the time point of the TF binding event, or the time point at which a motif transitions from being bound to unbound (lose events, E) or vice versa (gain events, D). Related to Figure S4.

We plotted the distribution of methylation across all timepoints for all binding events observed at each timepoint (**Figure 4D,E**). Overall, TF binding sites are both hypomethylated and accessible (**Figure S4B, S4C**); however, 30% of all accessible regions undergo some type of transition over the 12-day time course, both at the level of TF binding and DNAme. For all sites that lose TF binding at any timepoint, we find that hypomethylation is maintained long after binding sites are lost (**Figure 4D**). These data suggest that hypomethylation is intentionally maintained regardless of TF binding and accessibility. Furthermore, the transcriptional silencing of these regions cannot be attributed to the gain of DNAme, as transcription of neighboring genes closely follows TF binding activities. By contrast, for regions that gain a TF binding site at any timepoint during NPC differentiation, loss of DNAme begins to appear just prior to TF binding and, in general, this loss steadily continues after the binding event (**Figure 4E**). This was unexpected considering that TF binding is thought to be the initiator of demethylation and that resulting hypomethylation allows for stable TF binding. Overall, these data allowed us to resolve the order of events related to TF expression, binding and DNAme, revealing that demethylation activities start before appreciable TF binding is observed.

### Early and sustained accumulation of 5-hmC demarcates demethylation timing at lineage specifying enhancers

Of the three TET family members, TET1 and TET3 are highly expressed throughout the duration of our time course, in line with previous studies.^66^ While TET2 is less abundant than TET1/3, it is significantly upregulated (p-value = 0.0143) along with its co-factor IDAX (CXXC4, p-value = 0.0464) around 48 hours into differentiation, coinciding with the onset of substantial demethylation (**Figure S5A**). Likewise, global levels of 5-hmC increase significantly during differentiation, peaking at 4.5 days and decreasing to near baseline levels by day 12 (**Figure 5A**, ANOVA p=0.0228, Tukey’s HSD 0/108 p=0.0114, 6/108 p = 0.05069). Given the specific timing of demethylation and its apparent decoupling from ChrAcc changes, we examined the relationship between 5-hmC and cell cycle dynamics, as replication rates also change during hESC differentiation. We combined BrdU labeling and 5-hmC staining in a single flow cytometry panel to evaluate relative per-cell 5-hmC levels at each cell cycle stage (**Figure S5B, S5C**). We reasoned that, if 5-hmC is diluted during DNA synthesis, then levels of 5-hmC would be highest in G0 and G1 cells and would decrease as new DNA is synthesized. However, at all timepoints, cells in G2 displayed the highest 5-hmC, followed by S phase cells. These results support that a continuous, active demethylation mechanism is resolving 5-hmC to cytosine, as 5-hmC tracks more closely with total DNA content (**Figure 5B**).

**Figure 5:**
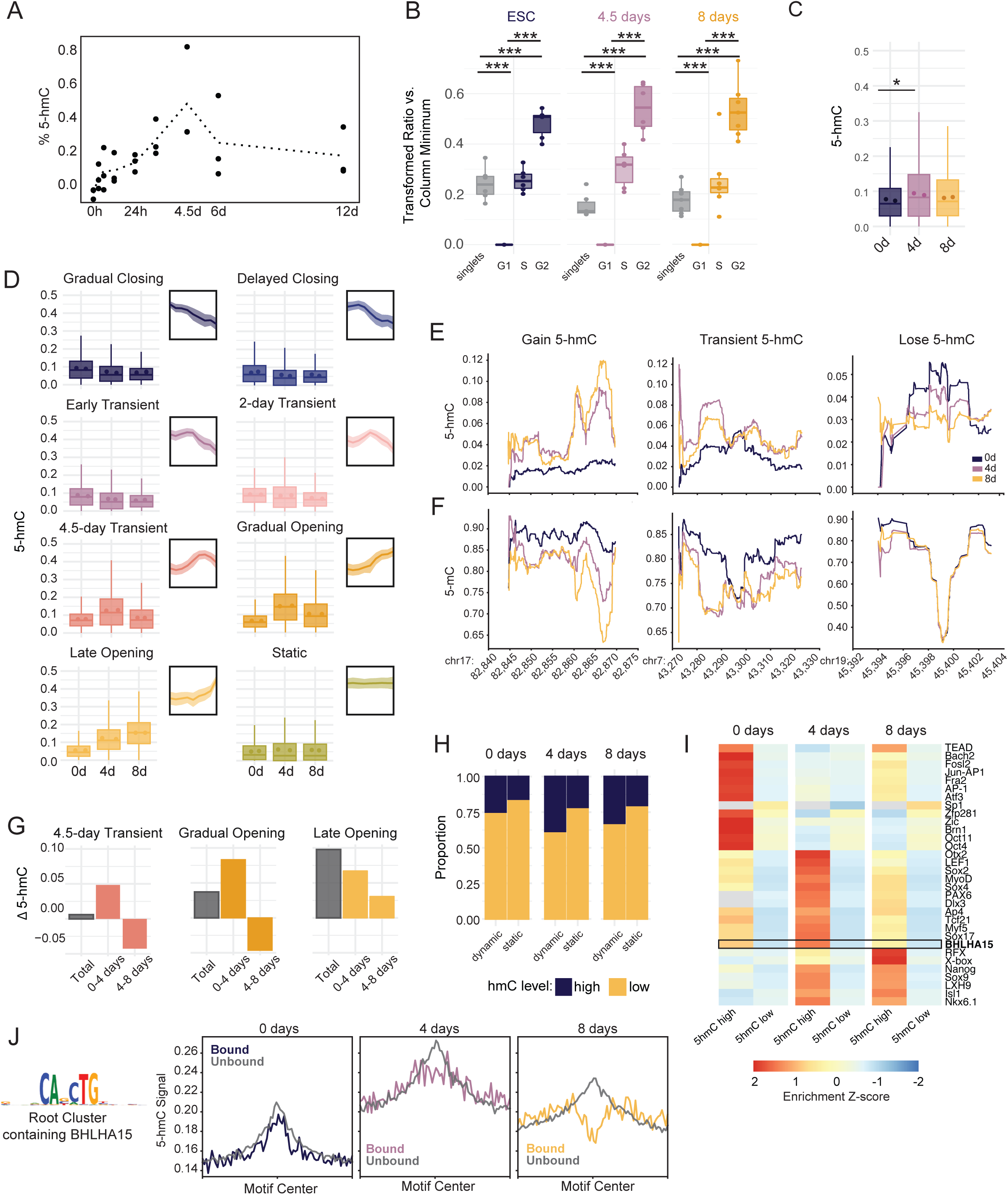
Early and sustained accumulation of 5-hmC demarcates demethylation timing at lineage specifying enhancers. (A) Dotted line plot shows the average global %5-hmC of biological replicates measured by ELISA at nine timepoints. Individual biological replicates are shown as black dots. Each biological replicate is the average of two technical replicates. % 5-hmC is determined via standard curve. (B) Boxplots display the distribution of 5-hmC signal across cell cycle stages for each timepoint. 5-hmC was measured by immunostaining and flow cytometry and is displayed as a transformed ratio versus the minimum median signal intensity using Cytobank.^98^ The transformed ratio was calculated using the minimum within each sample group (timepoint, See Methods). Events were gated into cell cycle stage using PI/BrdU staining, which is shown in **Figure S5B**. ANOVA and Tukey HSD were used to compare 5-hmC across cell cycle stages (p-value <2e-16 for all comparisons). (C) Boxplots show average proportion 5-hmC (reads reporting 5-hmC/total reads) at CpG sites within dynamic accessible peaks at 2, 4, and 8 days. 5-hmC proportion was measured using whole genome 6-base sequencing for two biological replicates. The mean proportion 5-hmC of individual replicates is shown for each timepoint as colored dots, *, p= 0.0365, one-sided t-test. (D) Boxplots display the average proportion 5-hmC (reads reporting 5-hmC/total reads) of CpG sites across regions in each accessibility cluster. Individual biological replicate means are displayed as points within the boxplot. Thumbnail visualizations of accessibility signal for each cluster are displayed. (E) Representative traces for proportion 5-hmC and (F) proportion 5-mC at three genomic loci displaying different types of 5-hmC changes between the three time points. Chromosome and coordinates (x1,000) for each locus are printed below the plot. Proportion 5-hmC is calculated as the average number of reads reporting 5-hmC over the average total number of reads for two biological replicates. Proportion 5-mC is calculated as the average number of reads reporting 5-mC over the average total number of reads for two biological replicates. CpGs with coverage less than 15 reads over both replicates were excluded for this analysis. (G) The average change in proportion 5-hmC was calculated for ChrAcc regions in three representative dynamic ChrAcc clusters. “Total” represents the average difference between 8-day and 0-day timepoints, “0-4 days” represents the difference between 4-day and 0-day timepoints, and “4-8 days” represents the difference between 8-day and 4-day timepoints. (H) The proportion of static and dynamic ChrAcc regions with high or low 5-hmC within at each 6-base timepoint. Regions with an average fraction 5-hmC ≥ 0.106 (top 25% of regional 5-hmC fractions) across replicates were termed “high” and regions with an average fraction 5-hmC < 0.106 across replicates were termed “low”. (I) Heatmap displaying motif enrichment for 5-hmC high and 5-hmC low regions at each timepoint. Motif enrichment is displayed as the fold-change over background and is scaled by TF across each row. Grey boxes represent values that were not significant (>0.05) at the respective timepoint. The boxed row represents the motif enrichment for BHLHA15 which is selectively enriched in regions with high 5-hmC at 4 days. (J) Aggregate profiles display 5-hmC signal at TF footprints for the JASPAR root cluster containing BHLHA15 (shown to the left). TF footprinting and binding state designation was performed using TOBIAS. Profiles display signal at footprint sites with a flanking distance of +/-1000bp. Signal is binned into 25bp bins. Related to Figure S5.

To quantify 5-hmC at nucleotide resolution, we performed 6-base sequencing, which is a whole genome sequencing approach capable of distinguishing between 5-mC and 5-hmC. We collected three timepoints in duplicate including 0 days, 4 days, and 8 days post-induction, as these timepoints capture the key phases of 5-hmC dynamics that we observed globally (**Figure 5A, 5C**). We quantified 5-hmC levels within our dynamic accessibility regions, finding that, unlike 5-mC, gain and loss of 5-hmC tracks closely with accessibility changes (**Figure 5D**). Example loci are depicted in **Figure 5E** to illustrate these trends at higher resolution. Moreover, 5-hmC levels increase prior to demethylation and then decrease as the demethylation process resolves, which is indicated by the decreased proportion of 5-mC in reads measured from the same locus (**Figure 5E, F**). This pattern is most clearly captured in *4.5 Transient* and *Gradual Opening* clusters, likely due to the timeframe when these regions are most accessible (**Figure 5E**). Regions that are open early show the highest level of 5-hmC at 0 days, prior to accessibility changes, but steadily decrease at 4 and 8 days (*Early Transient* and *2-day Transient*). In *4.5-day Transient*, *Gradual Opening*, and *Late Opening* groups, 5-hmC also increases prior to peak chromatin accessibility (**Figure S5D**). These regions display the greatest increase in 5-hmC between 0 and 4 days (**Figure 5G, S5D**). Closing regions display low levels of 5-hmC that decreases moderately over the time course, which supports the observation that closing regions continue to lose methylation even after returning to a closed state (**Figure 3D, S5E**). This means that demethylase activity begins early in the process to generate the 5-hmC levels that anticipate accessibility changes. 5-hmC also lingers as regions are returning to a closed state or as accessibility stabilizes, supporting the observation that complete loss of DNAme is delayed in regions that open.

Among dynamic regions, we observe a range of 5-hmC levels, indicating certain regions have greater 5-hmC than others (**Figure S5F**). We classified regions as “5-hmC high” if their regional average 5-hmC proportion was in the top 25% of all accessible regions. 5-hmC high regions were enriched within dynamic accessibility clusters compared to static regions, demonstrating a link between 5-hmC and ChrAcc dynamics (chi-squared: 0-day p-value < 2.2e-16, 4-day p-value < 2.2e-16, 8-day p-value < 2.2e-16, **Figure 5H**). We further observed that distinct subsets of TFs were specifically enriched in dynamic regions with high 5-hmC (**Figure 5I**). To examine 5-hmC and TF binding activity, we focused on dynamic regions with high 5-hmC at 4 days that contain BHLHA15 root motifs, which includes NeuroD2 (**Figure 5J, S5G**).^67^ While both bound and unbound sites display an accumulation of 5-hmC at 4 days, bound sites, but not unbound sites, displayed a dearth of 5-hmC in the region immediately surrounding the binding site which becomes more prominent by 8 days. This result, combined with the progressive loss of DNAme signal, suggests demethylase activity begins early, prior to TF binding, but that complete demethylation follows TF binding. These data raise the possibility that 5-hmC can forecast accessibility changes and TF binding, at critical enhancers prior to being resolved through demethylation.

### A machine learning approach predicts chromatin accessibility patterns from timepoint specific DNA methylation states

Previous machine learning approaches have used DNAme^68–71^, and more recently hydroxymethylation^72, 73^, to train models that predict gene expression or disease state. We developed a machine learning approach to test whether timepoint specific DNAme states can be used to predict past, present and future chromatin accessibility. Using XGBoost^74–76^, we began by training models separately on 5-mC, 5-hmC, and 5-mC + 5-hmC measured using 6-base sequencing (0, 4, and 8 days) for either dynamic or static ChrAcc regions. Timepoints were matched to their nearest temporal neighbor, such that predicted ChrAcc values from models trained on 0-, 4-, and 8-day methylation data were compared with observed ChrAcc values from 0, 4.5, and 12 days, respectively (**Figure S6A**). We tested each timepoint specific model on itself as well as other timepoints, generating a total of 9 models and 27 tests comparing observed vs. predicted ChrAcc (**Figure S6B**). For comparison, we also trained models on ChrAcc of enhancer and promoter regions using ENSEMBL annotations for NPCs or ESCs, irrespective of accessibility trend. Promoter trained models performed better at predicting promoter accessibility than those trained and tested on enhancers, with each timepoint performing equally well, especially when using models trained on both 5-mC and 5-hmC (**Figure S6C**). Similarly, we observed that models trained and tested on static ChrAcc regions performed better, on average, than models trained on dynamic regions (**Figure 6A, B, S6D**). In fact, static region models performed well at all timepoints, regardless of their training dataset (Spearman ρ>0.7). This is not surprising considering the prevalence of CpG dense promoter regions and other CpG islands in static regions, which are predominantly constitutively hypomethylated; thus, stable methylation states are highly predictive of stable ChrAcc states.

**Figure 6:**
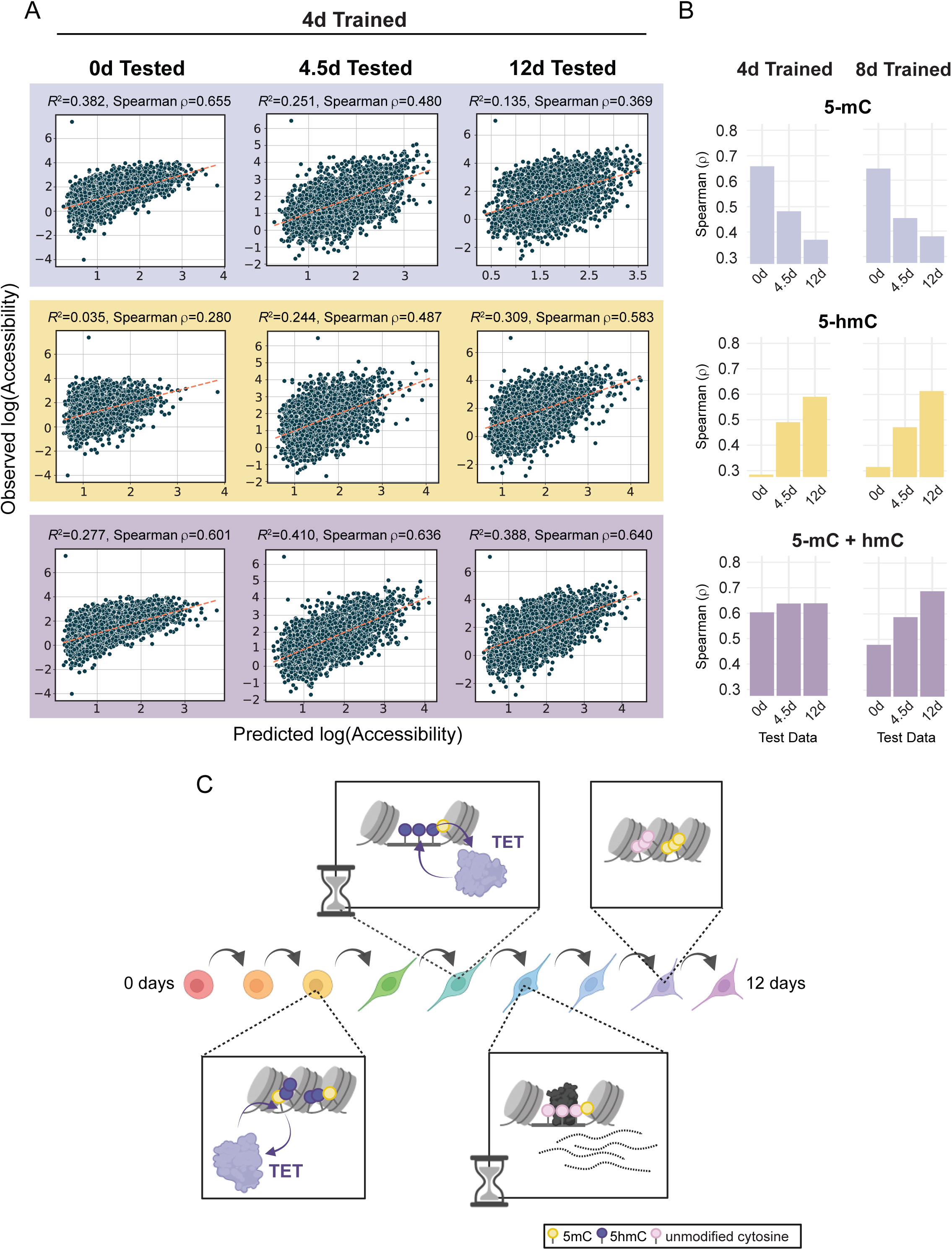
Chromatin accessibility prediction by machine learning. (A) Scatter plots display the observed accessibility versus the predicted accessibility for machine learning models trained on 4-day 5-hmC and 5-mC data (5-mC alone, 5-hmC alone, and 5-mC + 5-hmC). XGBoost models were trained on dynamic ChrAcc regions (excluding regions on chromosome 1) using methylation data from each singular timepoint (0, 4, and 8 days) and tested on regions from chromosome 1 at each timepoint. The spearman correlation coefficient is shown for each model. Dotted lines are defined by the slope between the points [minimum predicted value, minimum predicted value] and [maximum predicted value, maximum predicted value] in each scatterplot. (B) Bar plots of spearman ρ values (predicted vs. observed accessibility) for dynamic accessibility region models trained on 4-day or 8-day trained methylation data. Models were tested on all three timepoints in a similar fashion to those in A. Plots are divided by which methylation states were used for fitting. (C) A representative schematic of the molecular timeline proposed in this study. During the cell fate transitions that accompany NPC differentiation, enhancer regions that will be opened and activated first undergo 5-mC oxidation whereby 5-mC becomes 5-hmC (purple lollipops). This is followed by increases in accessibility and further oxidation, resulting in subsequent demethylation. TFs can bind these hydroxymethylated sites and facilitate the completion of demethylation while activating transcription of associated genes. Both the initial demethylation steps and the completion of the demethylation cycle are discretely timed events that occur between 2-6 days of differentiation. When an enhancer region is no longer required by the new cell fate, it loses TF binding and decreases in accessibility. However, the regions remain hypomethylated. Related to Figure S6.

To understand whether DNAme can predict ChrAcc in dynamic regions, we focused on models trained on 4-day methylation data (**Figure 6A, B**), which represents the timepoint for which 5-hmC was most frequently observed and coincides with the regions experiencing the greatest demethylation. While models trained on a combination of 5-mC and 5-hmC generally performed best at predicting ChrAcc, 5-mC and 5-hmC contributed differently to the model’s strength. For example, models trained on 5-mC alone performed best when tested at 0 days. This is especially true for 0-day trained data (**Figure S6E, F**). The strong model performance seen with ‘5-mC only’ models (compared to ‘5-hmC only’) tested on 0-day accessibility is likely due to the shortage of 5-hmC at 0 days, not to mention that most open chromatin regions are stably hypomethylated in HESCs. As expected, 0-day trained data performed poorly at predicting ChrAcc at 4 and 12 days.

By contrast, 4-day 5-mC + 5-hmC predictions showed higher correlations with observed accessibility levels at 0, 4 and 12 days (**Figure 6A**). Moreover, predictions from 5-hmC only models showed increasing correlation with observed accessibility from 0 to 12 days, indicating that 5-hmC contributes substantially to the 5-mC + 5-hmC models at later timepoints (**Figure 6B**). These performance trends are replicated in the 8-day trained models, which performed best at predicting accessibility at 12 days. It is also important to note that models trained on dynamic regions, the majority of which are lineage-specifying enhancers, performed substantially better at predicting dynamic accessibility than models trained and tested on enhancer annotations (**Figure 6A, S6C**). Overall, these results argue that, in order to understand the relationship between DNAme and ChrAcc and their joint role in regulating transcription, consideration of time and a combination of DNAme states is crucial (**Figure 6C**). By capturing this information, our data support the hypothesis that DNAme states can predict past, present and future chromatin states.

## DISCUSSION

Enhancers are activated progressively through recruitment of TFs and chromatin modifiers to permit access to DNA. Until recently, DNA demethylation was considered intrinsic to this process and essential for subsequent gene expression. However, in previous work we observed negligible enhancer demethylation during terminal cell differentiation despite robust ChrAcc and transcriptional changes.^1^ Similarly, steady state ChrAcc and DNAme data has previously revealed that accessible enhancers can be nucleosome free while also displaying a range of DNAme levels, including hypermethylation.^19, 20^ Further, the presence of DNAme at enhancers does not necessarily restrict TF binding or transcription of associated genes.^1, 2, 12, 45, 77^ While these observations challenge textbook models of DNAme and its role in gene regulation, how these discordant patterns are produced and their functional significance remains unclear.

In the present study, we address several important questions raised by previous work: First, our previous data was generated in cells that become post-mitotic, and the ability to observe substantial demethylation may be replication dependent.^49, 53, 78^ Here, we capture significant, primarily unidirectional, DNAme changes in proliferating NPCs over a substantially longer time course. Nonetheless, the decoupling of DNAme changes from ChrAcc and transcription still holds true, so the discordance between chromatin and DNAme changes is not a result of proliferative or developmental state.

Second, past studies did not distinguish 5-mC from 5-hmC, so the initiation or completion of demethylation could not be pinpointed relative to ChrAcc. Using densely sampled ATAC-Me data with 6-base sequencing, we show that, as enhancers experience waves of ChrAcc and TF binding, 5-hmC appears early but resolves late in the process. This temporal separation produces discordant epigenetic states at individual timepoints. In light of these new insights, the conclusion that enhancers are wholly insensitive to methylation may require some reconsideration, as enhancers that are both accessible and methylated may be under transition.

In addition, structural studies have demonstrated that TET1/2 are more efficient at catalyzing 5-mC than 5-hmC substrates, so complete removal of 5-hmC may take longer to resolve than the initial oxidation step.^79, 80^ This may explain, in part, why treatment with vitamin C, which enhances TET catalytic activity, increases DNAme loss in both mitotic and post-mitotic cells.^1, 81,82^ Indeed, non-physiological levels of vitamin C may accelerate the resolution of oxidized 5-mC substrates, which are not distinguished from 5-mC in bisulfite sequencing data. Alternatively, conversion of 5-mC to 5-hmC alone may be sufficient to permit transcription and TF binding rendering complete demethylation unnecessary. 5-hmC signal described here may also indicate an additional function outside of its role as a methyl-intermediate.^31^

While many TFs are considered insensitive to DNAme^20, 35, 36, 38–40^, their binding sites do ultimately display low DNAme levels, which we similarly observed. We examined DNAme levels from accessible DNA fragments before, during, and after predicted TF binding events. Loss of methylation appeared prior to TF binding and was corroborated by the presence of 5-hmC, which accumulated locally and diminished by subsequent timepoints. These findings indicate that the start of demethylation is at least concomitant with the start of TF binding. One caveat of our approach is that TF binding is indirectly determined by Tn5 cut-site frequencies, which is dependent on ATAC-Me sequencing depth. However, by integrating TOBIAS footprints with ChIP-seq data, we have previously shown that this method accurately distinguishes bound and unbound sites for specific TFs.^83^ Future studies may directly probe binding of TFs through ChIP-based methods, combined with DNAme quantification^84–87^, to better understand temporal relationships between TF binding and DNAme.

In proliferating cells, enhancer demethylation is likely achieved through a combination of TET-mediated active and replication-mediated passive mechanisms. ^46, 49, 53, 88^ Across nine timepoints over twelve days, we found a distinct window during which the greatest loss of DNAme occurs, coinciding with increased TET2 expression and peak 5-hmC levels. We found that the specific timing of demethylation could be not explained by replication dynamics, as 5-hmC levels track with DNA content, suggesting 5-hmC is not diluted passively in this system. A recent study combining metabolic labeling of DNA with mass spectrometry revealed that 5-hmC accumulates on parental single-stranded DNA post replication, which may support our conclusion that a continuous, active demethylation mechanism is resolving 5-hmC to cytosine^46^; however, we cannot concretely determine whether the resolution mechanism is base excision repair as observed in post-mitotic neurons.^50^ Regardless, the timing of DNA demethylation does not appear to be a result of changes in cell cycle dynamics.

Apart from losing DNAme, few ChrAcc regions gained methylation. This predominate loss of methylation was observed in both opening and closing regions and persisted throughout the time course. Previous studies found that patterns of DNA hypomethylation capture both active and historically active enhancers, and that hypomethylated regions accumulate as cells differentiate.^10, 17–19, 89, 90^ However, these studies lacked the temporal resolution to determine how hypomethylated regions are established and their relationship to ChrAcc. Our findings corroborate these studies and additionally demonstrate that transcriptional silencing does not require the acquisition of DNAme at enhancers of associated genes. For these decommissioned enhancers, what maintains the long-term hypomethylation state is unclear, but we speculate that it could be repressive TFs capable of binding nucleosomal DNA^91^, the exclusion of methyltransferases, or both.

Our studies uncover not only that 5-mC patterns reflect historical enhancer accessibility, but unexpectedly that 5-hmC can predict future accessibility. This stems from the finding that 5-hmC accumulates ahead of increasing accessibility at some sites. 5-hmC has been associated with dynamic enhancers and ChrAcc regions^92–96^, but our detailed temporal analysis of these epigenetic states allowed us to build a machine learning model that captures and predicts the relationship between 5-mC, 5-hmC, and ChrAcc. This work underscores the distinct and time-dependent relationship between these epigenetic features, which could be expanded upon to build models that are generalizable to differentiation-dependent accessibility changes across cellular systems.^72^ Ultimately, when considering the question of whether DNAme is deterministic of transcriptional patterns, our work argues that applying a comprehensive view of demethylation as a process, involving multiple intermediate states, is critical when evaluating the regulatory impact of DNAme.

## ACKNOWLEDGMENTS

We thank current and former members of the Hodges lab for helpful feedback and valuable critiques of the manuscript. We also thank Bruce Carter, John Karijolich, and Bill Tansey for their insights and discussions. Illustrations were made with Biorender.com. We are grateful for support of the project by NIH awards (R01 GM147078 to E.H., R01NS118580 to R.A.I., R01DK103831 and U54CA274367 to K.S.L.), Department of Defense Idea award (W81XWH-20-1-0522 to E.H.), an American Cancer Society (ACS) Institutional Research Grant (#IRG-15-169-56 to E.H.), the Ben & Catherine Ivy Foundation (to R.A.I), a gift from the Michael David Greene Brain Cancer Fund at the Vanderbilt–Ingram Cancer Center (to R.A.I), the Vanderbilt University Stanley Cohen Innovation Fund (to E.H), the VU School of Medicine Dean’s Faculty Fellow Award (to E.H) and funds from the Vanderbilt Ingram Cancer Center.

## AUTHOR CONTRIBUTIONS

Conceptualization, L.N.G., E.H.; Methodology, L.N.G., T.J.S., E.H.; Writing, L.N.G., T.J.S., E.H.; Formal analysis, L.N.G., T.J.S., Y.Y., A.J., R.I., E.H.; Investigation L.N.G., T.J.S., A.J.S., J.A.Y., E.H.; Supervision, Funding Acquisition and Resources, E.H., K.S.L., R.A.I., F.P.

## DECLARATION OF GENERATIVE AI AND AI-ASSISTED TECHNOLOGIES

Generative AI and AI-assisted technologies were not used in the preparation of this manuscript.

## SUPPLEMENTAL INFORMATION TITLES AND LEGENDS

Document S1. Figures S1-S6 and legends.

Document S2. Word Document containing Tables S1-S4.

## STAR METHODS

### Resource Availability

#### Lead contact

Further information and requests for resources and reagents should be directed to and will be fulfilled by the lead contact, Emily Hodges.

#### Materials Availability

All unique/stable reagents generated in this study are available from the lead contact without restriction.

#### Data and Code Availability

- ATAC-Me-seq, RNA-seq, single cell RNA-seq, and 6-base data have been deposited in the Gene Expression Omnibus (GEO) and are publicly available as of the date of publication. Accession numbers are listed in the key resources table.
- All code has been deposited in a publicly available GitHub Repository. Links to repositories are listed in the key resources table.
- Data can be visualized using the UCSC Genome Browser at the link listed in the key resource table.
- Any additional information required to reanalyze the data reported in this paper is available from the lead contact upon request.

### Experimental Model and Subject Details

#### Cell Culture and Treatments

H9 human embryonic stem cells (gift of Dr. Vivian Gamma, Vanderbilt University) were cultured in mTeSR1 (StemCell Technologies). Culture conditions were maintained at 5% CO_2_, 37°C and 80% humidity. During routine culture, H9 ESCs were maintained in colonies with daily media changes. Cells were passaged when 80% confluent, or approximately every 4-5 days using ReLeSR (StemCell Technologies).

#### Neural Progenitor Cell Differentiation

Neural progenitor cell differentiation was performed using the STEMdiff™ SMADi Neural Induction Kit, per the manufacturer’s instructions. Briefly, H9 ESCs were maintained as usual until 80% confluent. Cells were then dissociated using Accutase (StemCell Technologies) to generate a single cell suspension. Cells were pelleted and resuspended in Neural Induction Media with Y-27632 (StemCell Technologies) to a final concentration of 1x10^6^ cells/ml. Media was replaced daily for the next 5 days before being passaged again on day 6 of differentiation. On day 6, cells were similarly dissociated with Accutase (StemCell Technologies) to generate a single cell suspension. Cells were split 1 to 6 and plated into NIM with Y-27632 for the first 24 hours after plating. Cells were cultured for another 6 days before the final collection at 12 days of differentiation.

#### ATAC-Me

The ATAC-Me protocol used in this system was optimized and detailed previously^56^. Briefly, cells were harvested using Accutase (StemCell Technologies) and a single cell suspension was generated. Following collection, 200,000 cells were lysed, and nuclei were collected. Cells were pelleted by centrifugation and resuspended in a gentle lysis buffer to isolate nuclei. Nuclei were then incubated in Tn5 transposition reaction buffer with Tn5 assembled with methylated adaptors. Accessible DNA fragments underwent purification, oligo replacement, and gap repair. Fragments then undergo heat denaturation and sodium bisulfite conversion using the EZ-Methylation Gold Kit (Zymo). Libraries were amplified and indexed using 8-12 cycles of PCR. ATAC-Me libraries were sequenced using 2x150bp paired-end reads on the NovaSeq6000 instrument.

#### RNA-seq

RNA was collected from 1x10^6^ cells for each NPC differentiation time point by pelleting cells at 4°C, 500 R.C.F for 5 minutes. After removal of supernatant, cell pellet was resuspended in 1mL of TRIzol Reagent by repeatedly pipetting up/down with a 1mL micropipette tip. RNA was purified from Trizol according to manufacturer instructions. RNA-seq libraries were prepared using the NEBNext^®^ Ultra^™^ II RNA Library Prep according to manufacturer’s instructions. RNA-seq libraries were sequenced using 2x150bp paired-end reads on the NovaSeq6000 instrument.

#### scRNA-seq

Cells were prepared using a Papain Dissociation kit (Worthington Biochemical Corporation) according to the manufacturers protocol with some modification. Samples for sequencing were grown as previously described in a 6-well plate. Briefly, 2.5 mL of Papain + DNase solution was added to each well of a 6-well plate. Plates were shaken at 70 RPM at 37°C and 5% CO_2_ for 30 min. After incubation, cells were dissociated by pipetting up and down using a 1000μL pipette. Cells were incubated again under the same conditions for 10 more minutes prior to gentle pipetting with a 10mL pipette. Resulting cell suspension was transferred to a 15mL conical tube containing 5mL Earle’s medium + 3mL reconstituted inhibitor solution. Tube is inverted 3-5 times to mix. Cells are centrifuged at 300 x g for 7 minutes and supernatant is aspirated before resuspension of cells in 500μL 1x PBS. The PBS/cell suspension is then moved to a tube with a 35uM nylon mesh filter cap. Cells were encapsulated using a modified inDrop platform^99^, and sequencing libraries were prepared using the TruDrop protocol^100^. Libraries were sequenced in a S4 flow cell using a PE150 kit on an Illumina NovaSeq 6000^101, 102^.

#### Duet evoC 6-base Sequencing

Cells were collected at 0, 4, and 8 days after induction of differentiation using Accutase. Genomic DNA was collected and purified using phenol-chloroform extraction prior to being sonicated for 45 seconds in a Diagenode One sonication device (Diagenode) generating fragments with an average size of 250bp. Libraries were made using the duet evoC kit (biomodal) with 50ng of fragmented DNA according to manufacturer’s instructions. Final libraries were sequenced using 2x150bp paired-end reads on the NovaSeq6000 instrument.

#### 5-hmC ELISA

Genomic DNA was collected and purified using phenol-chloroform extraction. DNA was sonicated for 45 seconds in a Diagenode One sonication device (Diagenode) generating 200-600bp fragments. 5-hmC quantification was performed using the Quest 5-hmC DNA ELISA Kit (Zymo) according to the manufacturer’s instructions using 20ng of fragmented DNA as input.

#### Cell Cycle and 5-hmC Flow Cytometry

Flow cytometry was performed as previously described with modifications^103^. Cells were treated with 20μM BrdU in mTeSR or NIM for 1 hour. Cells were then collected using Accutase (StemCell Technologies), washed once with PBS, and resuspended in methanol. Cells were incubated overnight in methanol at 4°C with rotation to fix. After centrifugation and removal of supernatant, cells were resuspended in 100mM Glycine in PBS and incubated for 20 min at 25°C. Cells were centrifuged, and supernatant was removed before resuspension in 0.1% (v/v) Triton-X in PBS. Cells were incubated at 25°C for 30 minutes. After centrifugation and removal of supernatant, cells were resuspended in washing solution (0.5% BSA and 0.5% Tween in PBS) and incubated for 30 min at 25°C. Cells were counted at this step and cell count was normalized between samples for staining. Between each staining step, cells were washed three times in washing solution. 5-hmC staining was done using 100μL of PBS with 1:100 anti-5-hmC (Active Motif) overnight at 4°C followed by secondary staining using 100μL of washing solution with 1:200 anti-rabbit IgG CF750 (Sigma) for 1 hour at room temperature. Following secondary staining, cells were resuspended in 100μL of 0.5% BSA in PBS. To each sample, 15μL of FITC-α-BrdU (BD Biosciences) was added and incubated for 1 hour at room temperature. Finally, cells were washed before being resuspended in 300μL PI solution (0.4μg/mL PI, 8ng/μL RNase A, 0.5% BSA in PBS), incubated for 30 min at 25°C, and moved to a round bottom test tube with a cell strainer cap (Falcon). Samples were run on a 5 laser Fortessa instrument with FlowJo. Analysis and visualization were performed using Cytobank and ggplot2^104^. Signal was quantified as the fold-change in per-cell 5-hmC median fluorescence intensity per sample compared to the lowest median signal for same experiment. The inverse hyperbolic sine (arcsinh) with a cofactor was used to compare samples as previously described^105^. The arcsinh median of intensity value × with cofactor c was calculated as arcsinhc(x) = ln(x/c + √((x/c)^2^ + 1)). The cofactor (c) is a fluorophore-specific correction for signal variance.

### Quantification and statistical analysis

#### Chromatin accessibility prediction by machine learning

Machine learning models were generated in *python* (v3.11.0) using the *scikit-learn* (v1.1.3) and *modality* (v0.10.0) packages. The models were fit to predict chromatin accessibility from three layers of methylation data values (modC, mC, and hmC). Chromatin accessibility values were generated from filtered bams, merged by replicate (*bigWigs*), and normalized by the length of the region. Methylation values were derived from the biomodal 6-base duet evoC data and represented ‘modC,’ ‘mC,’ ‘hmC,’ and ‘mC + hmC’ average values tiled across genomic regions. The amount of CpGs per region were also recorded for model input. In the comparison between dynamic and static regions, dynamically accessible chromatin peaks were grouped together into a single BED file for input. For the comparison of regulatory regions, ‘enhancers’ and ‘promoters’ were selected from an Ensembl genome annotation file downloaded from their FTP server (https://ftp.ensembl.org/pub/current_regulation/homo_sapiens/GRCh38/annotation/); promoters and enhancers were selected by matching strings (“promoter” and “enhancer,” respectively) in the third column. To standardize BED region size, we determined the central base pair for each region and extended these +/-250 bp. Chromatin accessibility and methylation was mapped over the 500 bp region. Methylation windows were tiled at 500 bp intervals beginning at -1000bp and ending at +1000bp, resulting in 5 windows. Mapping was performed with the *pyranges*.*intersect()* function. We used *xgb.XGBRegressor*() from the *xgboost* (v1.7.1) package to initialize a machine learning model. Training and testing data was split on chromosome 1, estimating a 90:10% split (∼90.37:9.63% split among all peaks) such that training data included chromosomes 2-22, X, and Y. Model parameters were optimized with *GridSearchCV()* through the parameter space: *n_estimators* - 100-600, 200; *max_depth* - 3-8, 2; *eta* - 0.01-0.05, 0.01; *subsample* - 0.2-0.6, 0.1; *colsample_bytree* - 0.8-1.0, 0.05. For optimization, models were trained and tested on 0- and 8-day data, revealing identical optimized parameters. For subsequent analyses, the following parameter values were used: *n_estimators* - 500; *max_depth* - 7; *eta* - .02; *subsample* - 0.5; *colsample_bytree* - 0.95. Model performance was measured by mean squared error, r^2^, Pearson’s r, and Spearman’s ρ values. Plots display Spearman’s ρ values and were generated in *ggplot2* (v3.3.6) in *R (v4.1.2)*.

#### ATAC-Me Library Processing

All ATAC-Me library reads were trimmed of adapters using TrimGalore script wrapper for Cutadapt^106^ and FastQC using the --fastqc and --paired parameters. ATAC-Me reads were mapped with WALT^107^ to the hg38 genome assembly using the -sam -m 6 parameters. Methylation analysis of ATAC-Me reads was performed using the MethPipe (v5.0.1, now DNMTools) suite of tools^108^. Symmetrical CpGs with 5 reads or greater coverage were included in all analyses. Proportion methylation at symmetrical GpGs were calculated using symmetric-cpgs from the MethPipe package with default settings after duplicates were removed. Mapped reads were filtered using samtools^109^ to exclude reads on ChrM, reads within blacklisted regions, and read with a MAPQ < 30. Regions enriched for chromatin accessibility in ATAC-Me data were identified using the Genrich (available at https://github.com/jsh58/Genrich) peak caller with the following parameters: -r -e chrX,chrY,chrM -j -p 0.005 -q 0.01 -v . Regions displaying dynamic chromatin accessibility were identified with the TCseq R-package^59^. Regional methylation levels were determined by roimethstat from MethPipe. HOMER was used for all transcription factor motif analysis of dynamic or static chromatin accessible regions without background. Annotation and gene association for dynamic and static chromatin accessible regions was performed with the ChIPseeker^110^ and ClusterProfiler^111^ R-packages. Transcription factor footprinting was performed on ATAC-Me libraries using the TOBIAS suite of tools^63^. The samtools^109^, bedtools^112^ and deeptools^113^ suites of tools were used to aid in data manipulation and visualization. Preseq^114^ was used to compare library complexity across timepoints for ATAC-Me libraries.

#### RNA-seq Library Processing

RNA libraries were mapped with the STAR aligner^115^ run on untrimmed reads using the following parameters: --runMode alignReads --runThreadN 8 --outSAMtype BAM SortedByCoordinate --quantMode GeneCounts. Mapped reads were filtered using samtools^109^ to exclude reads on ChrM, reads within blacklisted regions, and read with a MAPQ < 30. Read coverage across transcripts was determined through featurecounts^116^ using the Gencode v38 annotation file. Preseq^114^ was used to compare library complexity across timepoints for RNA-seq libraries. Differential RNA expression was performed using DESeq2^117^.

#### 6-base Library Processing

6-base sequencing libraries were analyzed with the duet pipeline (v1.2.0)^57^. Briefly, FASTQ files were trimmed and quality-filtered using cutadapt^118^, and the epigenetic states in each read pair were then resolved using couplet. Resolved reads were then aligned using BWA-MEM^119^ to a standard four-base reference genome comprising of both GRCh38 and spiked-in control sequences. Quantification of epigenetic modifications was calculated at each CpG context = present in the reference genome and covered in the sequencing. Further downstream processing was performed using the modality suite, developed by biomodal. For regional analyses, cytosines with a read coverage >= 15 over both replicates were included. modality (v0.10.0), bedtools^112^, and ggplot2 were used to aid in data manipulation and visualization.

#### scRNA-seq Library Processing

Single cell RNA-seq libraries were analyzed as done previously^101^. Briefly, reads were demultiplexed, aligned, and corrected with the DropEst pipeline^120^, using the STAR^115^ aligner with reference genome hg38 and paired with the corresponding GTF annotations. We identified high-quality, cell-containing droplets and their respective barcodes through a QC pipeline previously described^121^.

#### Quantification and Statistical Analysis

ATAC-Me chromatin accessibility peaks were filtered using the Benjamini-Hochberg corrected p value (q-value) reported by the Genrich peak-calling algorithm (corr. p value < 1x10^−10^). Differentially accessible genomic loci across the time course were selected using the TCseq R-package, utilizing a FDR corrected p value cutoff produced by the likelihood ratio test implemented in the R-package (corr. p value < 5x10^−3^). Differentially expressed genes were filtered using corrected p values produced by the likelihood ratio test implemented in the DESeq2 R-package for the comparison between the 0 day and 12-day timepoints (corr. p value < 5x10^−3^). Statistical analyses were performed within the R computing environment and visualized with ggplot2^104^ or deeptools^113^. Specific statistical analyses can be found in relevant figure legends. All visualization and analysis code can be found on our Github page.

