## Supplemental Materials for "Temporally discordant chromatin accessibility and DNA demethylation define short and long-term enhancer regulation during cell fate specification"

### SUPPLEMENTARY FIGURE LEGENDS

**Supplementary Figure 1: Directed differentiation of HESCs to NPCs displays extensive DNA demethylation within chromatin accessibility loci.** (A) Transcript expression as determined by RNA-seq is shown for canonical ESC and NPC markers. The displayed data represents the average *FPKM* normalized read counts for two biological replicates. (B) Library complexity was calculated using *preseq* for each sequencing library. The number of distinct reads for the sample is displayed. (C) Correlation between biological ATAC-Me replicates was calculated to evaluate reproducibility across replicates using *multiBigWigSummary*. The data points and Spearman correlation coefficient for each comparison is plotted. (D) Correlation between biological RNA-seq replicates was calculated to evaluate reproducibility across replicates using *multiBigWigSummary*. The data points and Spearman correlation coefficient for each comparison is plotted. (E) Static ChrAcc regions over the time course were identified using *TC-seq* (dynamic n = 38189, static n = 63026, $|\log_2\text{-fold}| > 2$, adjusted p-value < 0.05). *deeptools* is used to display the ChrAcc and DNAme values of these static peaks at all time points. Regions are sorted by decreasing normalized read count signal intensity at the 0-hour time points and are consistent across all heatmaps. Regions are scaled relative to each other along the center of each heatmap with the up and downstream flanking region (Methylation = +/-1kb, Accessibility = +/-0.5kb) . Related to Figure 1.

**Supplementary Figure 2: Unsupervised clustering of chromatin accessibility reveals temporally distinct regulatory groups with divergent changes in enhancer states.** (A) The *factoextra* package was used with the “wss”, “silhouette”, or “gap_stat” methods to calculate the total within-cluster sum of square, the average silhouette of observations, or the estimated gap statistic, respectively, for 1-20 clusters to inform the C-means cluster number for TC-seq. (B) Gene ontology enrichment was performed on genes associated with any dynamic cluster region using GREAT with the default, basal plus extension method of association. Ontology is grouped by dynamic cluster though there was no significant enrichment for regions contained in *Early Transient* and *2-day Transient* clusters (padj <0.05). The dot plot represents enrichments across biological processes, molecular function, and cellular component analyses. (C) Gene ontology enrichment was performed on genes associated with accessible regions using GREAT for biological processes. Ontology is grouped by accessibility behavior, static versus dynamic. The top 10 most significant processes are shown for dynamic regions, while all processes passing the significant threshold (padj <0.05) are displayed for static regions due to lower enrichment. (D) Annotation of regions by chromatin state was performed using the chromHMM[32] 18-state annotations from HESCs and NPCs. A Sankey plot displays the change in the regions' chromatin state from the ESC to NPC stages. *Gradual Opening* and *Late Opening* were consolidated for this visualization. (E) A Sankey plot displays the change in chromatin state for *Gradual Closing* and *Delayed Closing* which were consolidated for this visualization. (F) Histograms represent the distribution of observed/expected CpG ratio for regions contained in each accessibility cluster. The mean of all regions is included for all clusters. The ratio is calculated as follows: $Number\ of\ CpG * N\ /\ (Number\ of\ C\ *\ Number\ of\ G)$, where N = length of the region. (G) Average Obs/Exp CpG ratios were calculated for each major ChrAcc trend, with the distribution of regional values shown by violin plot. Mann Whitney U test was performed for individual comparisons to the *Transient* regions (*Transient vs. Closing* p-value = 4.507e-8, *Transient vs. Opening* p-value <2e-16, *Transient-Static* p-value <2e-16). Related to Figure 2.

**Supplementary Figure 3: DNAme dynamics are unidirectional and temporally discordant with chromatin accessibility.** (A) Line plot of the standardized difference of the mean across the time course for all dynamic loci, one standard deviation from the mean is shown by gray ribbon. Standardized difference of the mean across time was calculated for the average regional DNA methylation state of dynamic accessible fragments. (B) Boxplots display the change in average methylation values across regions between the first (0 day) and last (12 day) timepoints.

Regions are grouped by their accessibility behavior. (C) Violin plots display the distribution of regional methylation values for all accessible regions in each methylation cluster. Methylation clusters were defined by hierarchical clustering using the *ward.D2* clustering algorithm. 25,964 stably hypomethylated regions displaying <10% 5-mC across all time points are not displayed. (D) Scatterplots display methylation measurements of CpG sites within ChrAcc peak regions quantified by ATAC-Me or 6-base sequencing. (E) Boxplots display the standard difference of the mean across time for ChrAcc and gene expression. Standardized differenced was calculated using $\log_2$(normalized ATAC read counts +1) and $\log_2$(normalized read counts+1) of neighboring transcripts (top 25% most variable). Lines represent median values. Related to Figure 3.

**Supplemental Figure 4: Discordant DNAme patterns generate historical hypomethylation records.** (A) Normalized read counts for three JASPAR POU family root members, POU5F1, POU3F1, and POU3F2, derived from RNA-seq data captured at each timepoint. Data was normalized for sequencing depth using FPKM at each timepoint. (B) Average regional methylation and accessibility (C) for all TF binding sites that were called as bound or unbound by TOBIAS. Binding site regions included the TF motif sequence +/-50 bp. Accessibility signal is displayed as the $\log_2$ transformed ATAC signal. ATAC-Me read count was normalized using DESeq2 prior to transformation. (D, E) Line plots show average methylation values over time visualized by TF binding behavior. Methylation values are averaged across CpGs contained in the TF motif +/-50bp. The annotated time represents the time point of the TF binding event, or the time point at which a motif transitions from being bound to unbound (lose events, D) or vice versa (gain events, E). Shaded ribbons represent one standard deviation above or below the mean methylation. Related to Figure 4.

**Supplementary Figure 5: Early and sustained accumulation of 5-hmC demarcates demethylation timing at lineage specifying enhancers** (A) Heatmap displaying transcript levels for TET family members and the TET2 binding partner CXXC4/IDAX normalized using

FPKM. Values were $\log_2$ transformed prior to plotting and each transcript expression is scaled across the time course using the z-score. (B) Gating parameters were applied at each timepoint to capture single cell events within each stage of the cell cycle. Representative gating for one biological replicate is shown. Axes represent signal intensity and are labeled according to the channel being visualized. (C) Representative histograms of median fluorescent intensity for 5-hmC signal across cell cycle stages as measured by flow cytometry, produced in Cytobank. Histograms colors represent the transformed ratio relative to the minimum within the displayed representative sample. (D) Boxplots display the standard difference of the mean across time for DNA methylation, chromatin accessibility, and 5-hmC. Standardized differenced was calculated using $\log_2$(normalized ATAC read counts +1) the average fraction 5-mC of the chromatin accessible DNA fragments, and the  average fraction 5-hmC of accessible peak regions. Lines represent median values. 6-base data for three timepoints (0, 4 and 8 days) is displayed alongside ATAC-Me data for the timepoints closest to 6-base timepoints (0-day, 4.5-day, 6-day, and 12-day). (E) Change in mean 5-hmC proportion for accessibility clusters not displayed in Figure 5G. Regional means are averaged across all regions contained in the cluster. “Total” represents the difference between 8-day and 0-day data, “0-4 day” represents the difference between 4-day and 0-day data, and “4-8 day” represents the difference between 8-day and 4-day data. (F) The histogram displays the distribution of average regional 5-hmC fraction across all dynamic regions. (G) Aggregate profiles display 5-mC signal at TF footprints for a JASPAR root cluster containing BHLHA15. TF footprinting and binding state designation was performed using TOBIAS. Profiles display signal at footprint sites with a flanking distance of +/-1000bp, binned into 25bp bins. Related to Figure 5.

**Supplementary Figure 6: Chromatin accessibility prediction by machine learning.** (A) Diagram of timepoints sampled by ATAC-Me and 6-base methylation data assays used for machine learning models. ATAC was measured at 0, 4.5, and 12 days. 5-mC along with 5-hmC was assayed at 0, 4, and 8 days with 6-base sequencing. (B) Schematic of machine learning model workflow. A BED file representing enhancers, promoters, static accessibility regions, or dynamic accessibility regions was used as input for prediction targets. Training and testing datasets were split on chromosome 1. Training was performed on regions from all chromosomes withholding chr1, and testing was performed on chr1 regions. Input for training data consisted of methylation levels at individual CpG sites measuring 5-mC, 5-hmC, or both from 6-base sequencing data performed at 0, 4, or 8 days. Models were subsequently tested on ATAC data derived from 0, 4.5, or 12-day datasets. (C) Dot and line plots of spearman $\rho$ values of XGBoost models trained across the same three methylation data combinations. Models were similarly trained on the methylation of one day and tested across all three timepoints. Input data for models were subdivided into enhancers or promoter regions. Model performance by input methylation datatype is distinguished by line color. (D) Bar plots of spearman $\rho$ values (predicted vs. expected accessibility) for static region models trained on methylation values from 0, 4, or 8-day data and tested to predict accessibility at 0, 4.5, and 12 days. (E) Bar plots of spearman $\rho$ values (predicted vs. expected accessibility) for dynamic region models trained on 0-day data. (F) Scatter plots display the observed accessibility versus the predicted accessibility for models trained and tested on 0-day data across either dynamic or static accessibility regions. Dotted lines are defined by the slope between the points [minimum predicted value, minimum predicted value] and [maximum predicted value, maximum predicted value] in each scatterplot. Related to Figure 6.

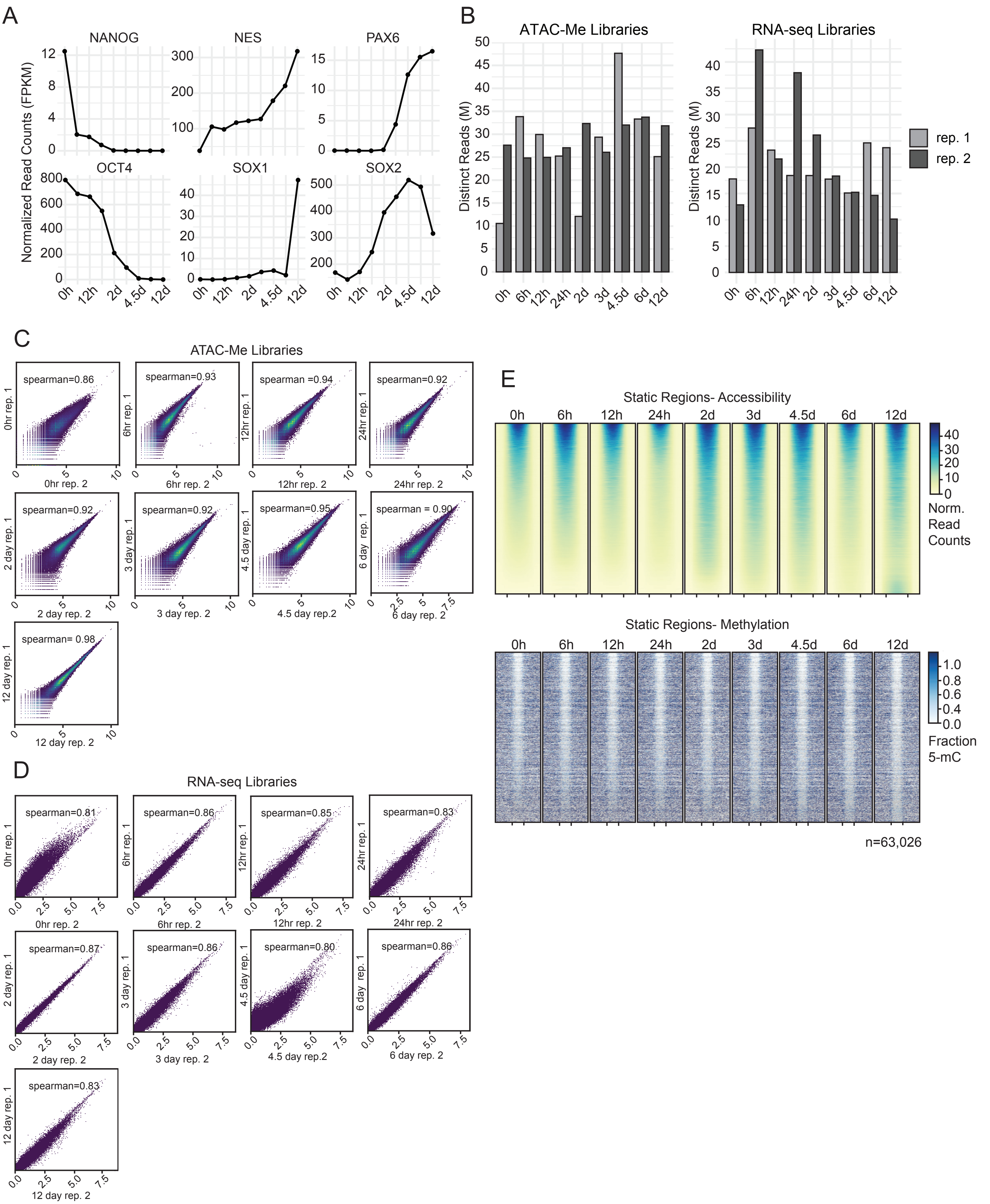
A
NANOG
NES
PAX6
OCT4
SOX1
SOX2
Normalized Read Counts (FPKM)
0h 12h 2d 4.5d 12d
B
ATAC-Me Libraries
RNA-seq Libraries
Distinct Reads (M)
rep. 1
rep. 2
0h 6h 12h 24h 2d 3d 4.5d 6d 12d
C
ATAC-Me Libraries
spearman=0.86
spearman=0.93
spearman =0.94
spearman=0.92
spearman=0.92
spearman=0.92
spearman=0.95
spearman = 0.90
spearman= 0.98
0hr rep. 1
0hr rep. 2
6hr rep. 1
6hr rep. 2
12hr rep. 1
12hr rep. 2
24hr rep. 1
24hr rep. 2
2 day rep. 1
2 day rep. 2
3 day rep. 1
3 day rep. 2
4.5 day rep. 1
4.5 day rep.2
6 day rep. 1
6 day rep. 2
12 day rep. 1
12 day rep. 2
D
RNA-seq Libraries
spearman=0.81
spearman=0.86
spearman=0.85
spearman=0.83
spearman=0.87
spearman=0.86
spearman=0.80
spearman=0.86
spearman=0.83
E
Static Regions- Accessibility
0h 6h 12h 24h 2d 3d 4.5d 6d 12d
Norm.
Read
Counts
Static Regions- Methylation
Fraction
5-mC
n=63,026

Guerin et al, Supplementary Figure 2

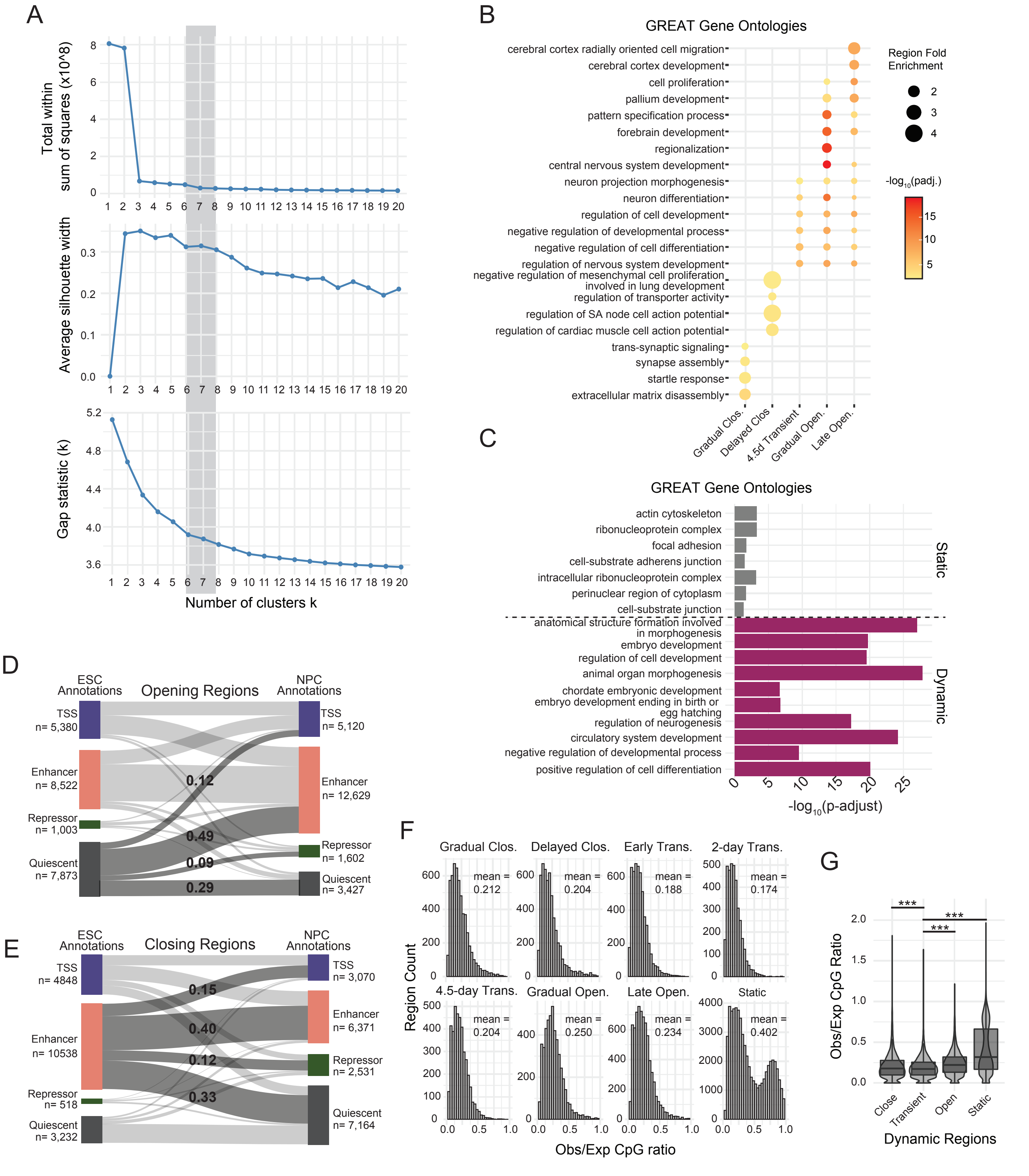

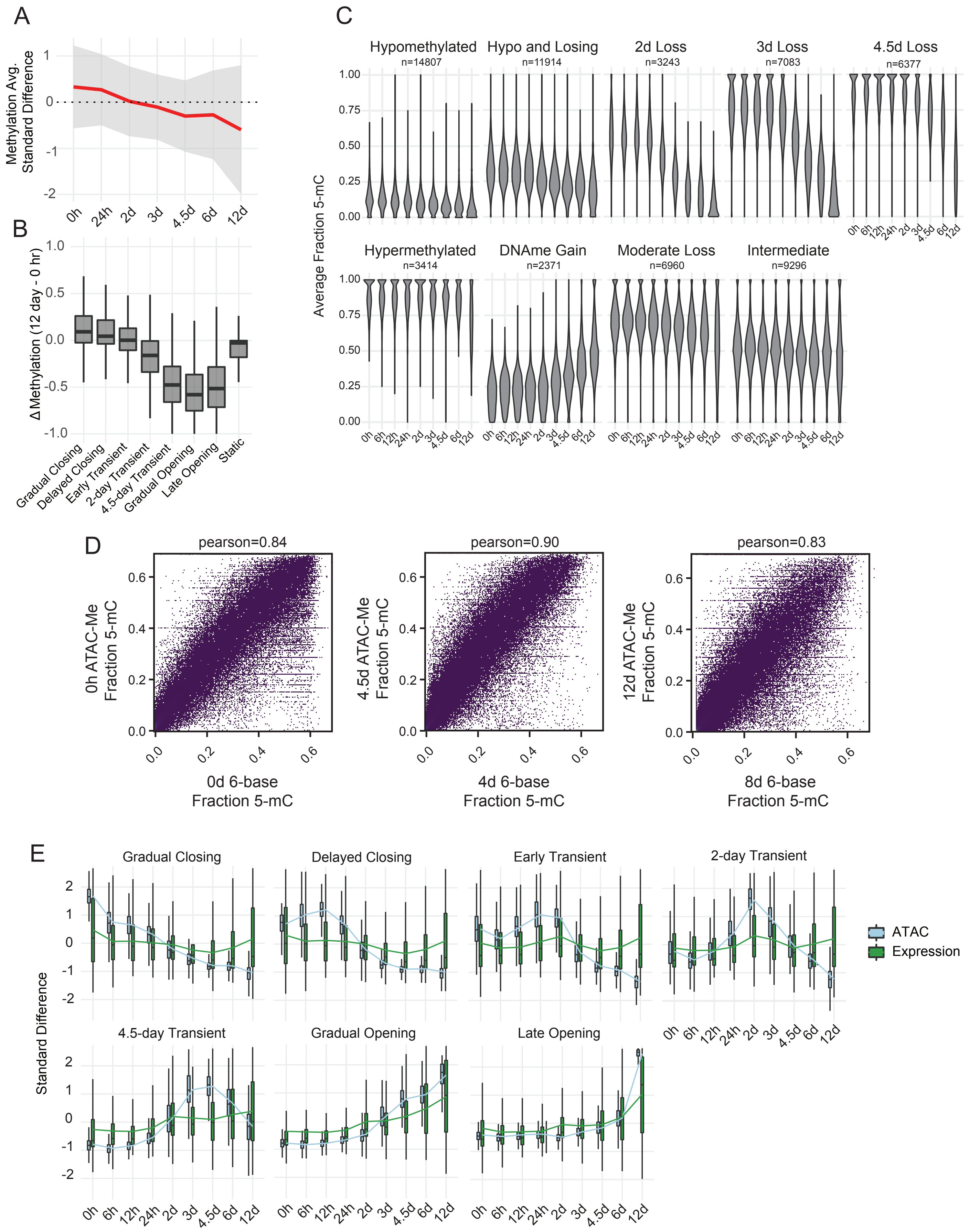
A
Methylation Avg. Standard Difference
0h
24h
2d
3d
4.5d
6d
12d
B
Δ Methylation (12 day - 0 hr)
Gradual Closing
Delayed Closing
Early Transient
2-day Transient
4.5-day Transient
Gradual Opening
Late Opening
Static
C
Average Fraction 5-mC
Hypomethylated
n=14807
Hypo and Losing
n=11914
2d Loss
n=3243
3d Loss
n=7083
4.5d Loss
n=6377
Hypermethylated
n=3414
DNAme Gain
n=2371
Moderate Loss
n=6960
Intermediate
n=9296
D
pearson=0.84
0h ATAC-Me Fraction 5-mC
0d 6-base Fraction 5-mC
pearson=0.90
4.5d ATAC-Me Fraction 5-mC
4d 6-base Fraction 5-mC
pearson=0.83
12d ATAC-Me Fraction 5-mC
8d 6-base Fraction 5-mC
E
Standard Difference
Gradual Closing
Delayed Closing
Early Transient
2-day Transient
4.5-day Transient
Gradual Opening
Late Opening
ATAC
Expression

Guerin et al, Supplementary Figure 4

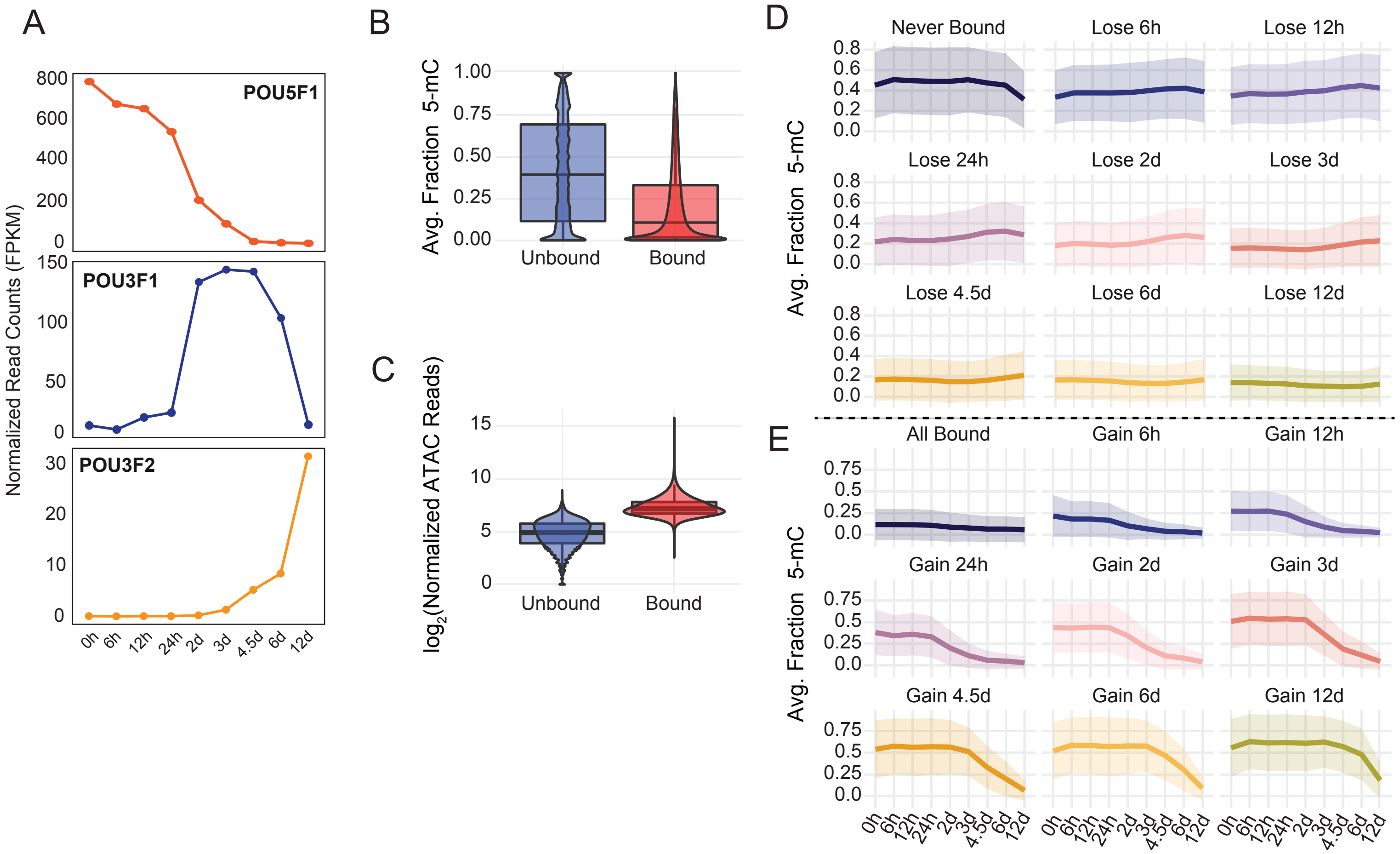

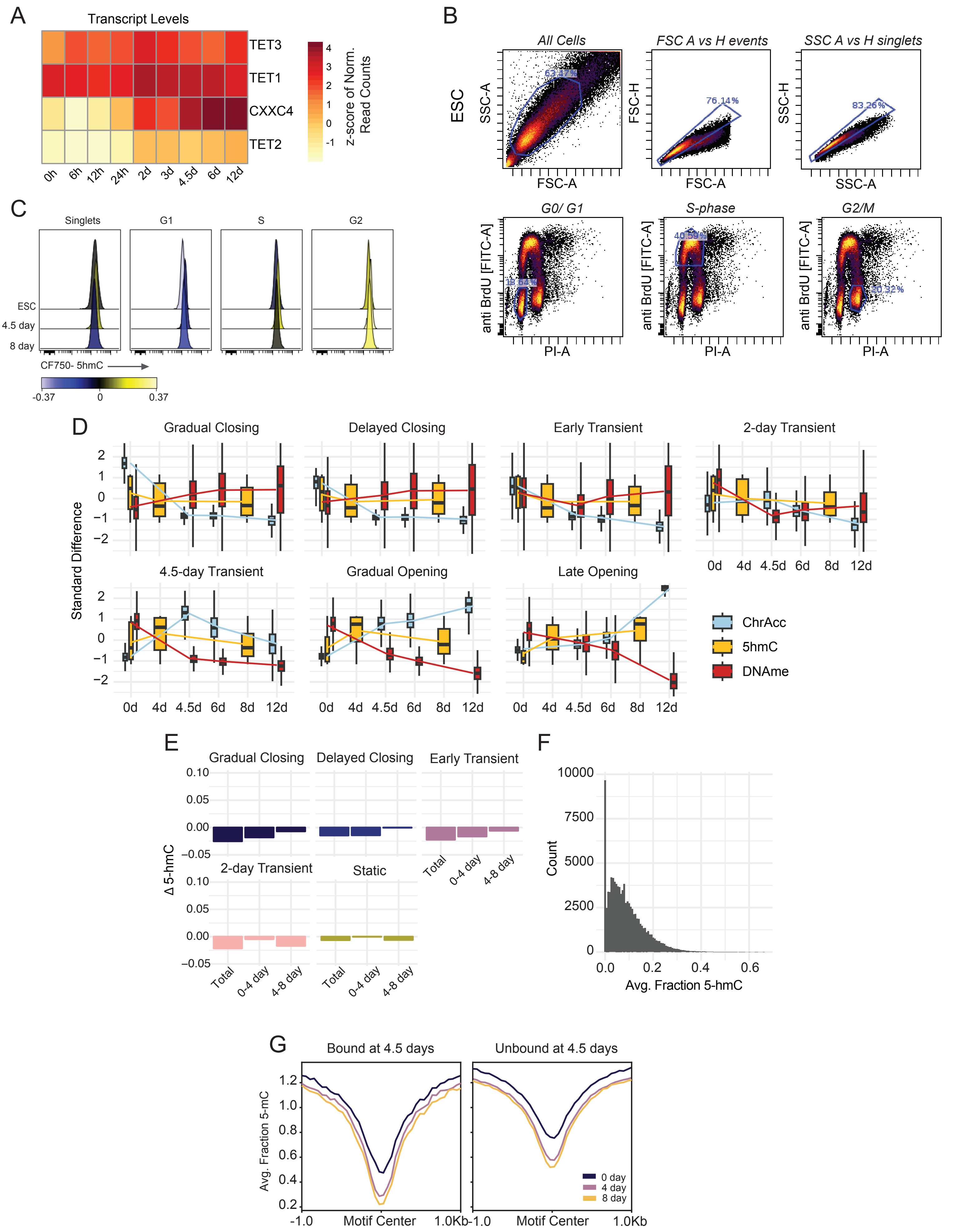
A
Transcript Levels
TET3
TET1
CXXC4
TET2
0h
6h
12h
24h
2d
3d
4.5d
6d
12d
z-score of Norm. Read Counts
B
ESC
All Cells
FSC A vs H events
SSC A vs H singlets
SSC-A
FSC-A
FSC-H
SSC-H
76.14%
83.26%
G0/ G1
S-phase
G2/M
anti BrdU [FITC-A]
PI-A
40.59%
20.32%
C
Singlets
G1
S
G2
ESC
4.5 day
8 day
CF750- 5hmC
-0.37
0
0.37
D
Gradual Closing
Delayed Closing
Early Transient
2-day Transient
4.5-day Transient
Gradual Opening
Late Opening
Standard Difference
0d
4d
4.5d
6d
8d
12d
ChrAcc
5hmC
DNAme
E
Gradual Closing
Delayed Closing
Early Transient
2-day Transient
Static
Δ 5-hmC
Total
0-4 day
4-8 day
F
Count
Avg. Fraction 5-hmC
G
Bound at 4.5 days
Unbound at 4.5 days
Avg. Fraction 5-mC
Motif Center
-1.0
1.0Kb
0 day
4 day
8 day

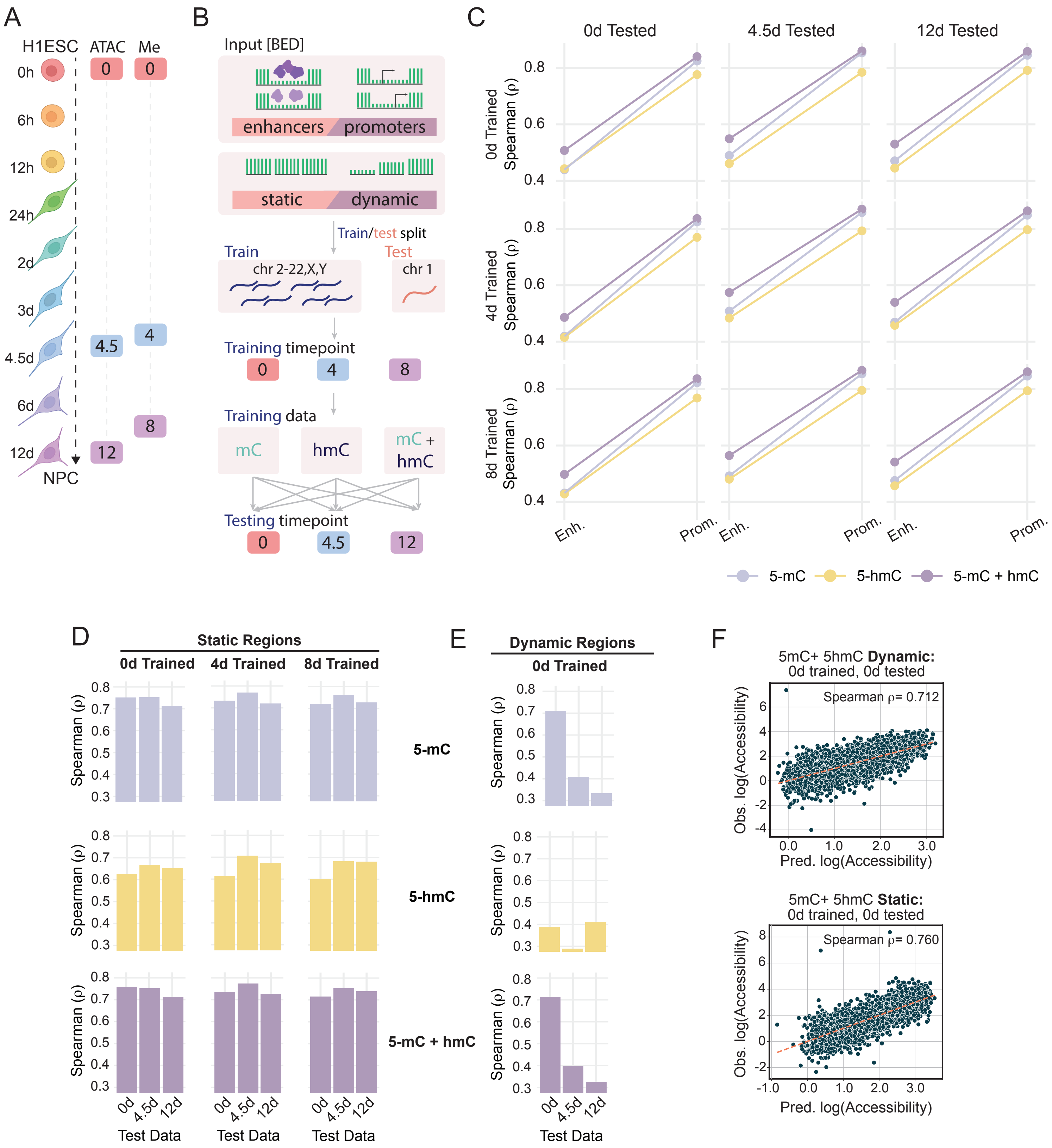
A
H1ESC
ATAC
Me
0h
6h
12h
24h
2d
3d
4.5d
6d
12d
NPC
B
Input [BED]
enhancers
promoters
static
dynamic
Train/test split
Train
Test
chr 2-22,X,Y
chr 1
Training timepoint
Training data
mC
hmC
mC + hmC
Testing timepoint
C
0d Tested
4.5d Tested
12d Tested
0d Trained Spearman (ρ)
4d Trained Spearman (ρ)
8d Trained Spearman (ρ)
Enh.
Prom.
5-mC
5-hmC
5-mC + hmC
D
Static Regions
0d Trained
4d Trained
8d Trained
Spearman (ρ)
5-mC
5-hmC
5-mC + hmC
Test Data
E
Dynamic Regions
0d Trained
F
5mC+ 5hmC Dynamic: 0d trained, 0d tested
Spearman ρ= 0.712
Obs. log(Accessibility)
Pred. log(Accessibility)
5mC+ 5hmC Static: 0d trained, 0d tested
Spearman ρ= 0.760

**Table S1:** Library, sequencing, and analysis statistics for each ATAC-Me sample.

| ATAC-Me Sequencing | | | | |
|---|---|---|---|---|
| Sample | Final library concentration (ng/ul) | Total reads | CpGs (coverage ≥ 5X) | Peak number |
| 0hr A | 5.93 | 67213113 | 1302926 | 46961 |
| 0hr B | 1.09 | 58626341 | 2089056 | |
| 6hr A | 6.88 | 103292803 | 2981980 | 46915 |
| 6hr B | 16 | 60019065 | 2036771 | |
| 12hr A | 11.1 | 88052404 | 2650643 | 42019 |
| 12hr B | 5.88 | 68990381 | 1732449 | |
| 24hr A | 6.86 | 68152510 | 2124207 | 27765 |
| 24hr B | 5.32 | 69078947 | 1869887 | |
| 48hr A | 1.05 | 177315084 | 2079575 | 65270 |
| 48hr B | 8.28 | 79665573 | 2696196 | |
| 72hr A | 15.3 | 90632062 | 3478202 | 57044 |
| 72hr B | 2.71 | 71355801 | 2347331 | |
| 4.5day A | 2.23 | 105550194 | 4217114 | 58888 |
| 4.5day B | 8.16 | 72500021 | 2769749 | |
| 6day A | 4.36 | 55513844 | 2995283 | 33418 |
| 6day B | 9.65 | 60194982 | 2684612 | |
| 12day A | 13.4 | 55598687 | 2067362 | 59581 |
| 12day B | 8.54 | 75398649 | 2456311 | |

**Table S2:** Library, sequencing, and analysis statistics for each RNA-seq sample.

| RNA sequencing | | | | |
|---|---|---|---|---|
| Sample | Final library concentration (ng/ul) | Input reads | Uniquely mapped reads | Filtered reads |
| 0hr A | 27.8 | 31909040 | 22158598 | 8610268 |
| 0hr B | 20.3 | 27715446 | 18288951 | 6593643 |
| 6hr A | 0.684 | 49931050 | 35668744 | 13271468 |
| 6hr B | 5.27 | 16216974 | 12303359 | 20646873 |
| 12hr A | 22 | 43495053 | 29803224 | 11368648 |
| 12hr B | 19.1 | 23324620 | 17976459 | 10318180 |
| 24hr A | 15.3 | 32461243 | 23247257 | 8797646 |
| 24hr B | 24.9 | 25032726 | 18718534 | 17894918 |
| 48hr A | 17.1 | 30598220 | 22546414 | 8750219 |
| 48hr B | 17.5 | 44161371 | 32901592 | 12379472 |
| 72hr A | 6.66 | 34465624 | 22359612 | 8632510 |
| 72hr B | 30.9 | 31959301 | 22174802 | 8709773 |
| 4.5day A | 7.25 | 31953740 | 19735284 | 8013688 |
| 4.5day B | 20.3 | 63603786 | 48041193 | 7378077 |
| 6day A | 4.38 | 49554234 | 32078907 | 12392128 |
| 6day B | 15.8 | 36253322 | 27096703 | 7138882 |
| 12day A | 17.8 | 41643300 | 28975427 | 11522785 |
| 12day B | 12.7 | 76323733 | 57066247 | 4910512 |

**Table S3:** Full list of TF motifs enriched in each cluster, as visualized in Figure 2D.

| log_pval | percentFold | Motif | Cluster |
|---|---|---|---|
| -2031 | 9.99126638 | OCT4-SOX2-TCF-NANOG | Delayed Closing |
| -1111 | 3.9653092 | OCT4 | Delayed Closing |
| -927.5 | 3.98214286 | OCT6 | Delayed Closing |
| -879.1 | 4.59305211 | Brn1 | Delayed Closing |
| -834.8 | 4.89675516 | Jun-AP1 | Delayed Closing |
| -812.1 | 4.10042735 | Fosl2 | Delayed Closing |
| -804.8 | 3.43647235 | Fra2 | Delayed Closing |
| -767.4 | 3.15374677 | Fra1 | Delayed Closing |
| -695 | 2.76012461 | Atf3 | Delayed Closing |
| -688.9 | 2.98867925 | JunB | Delayed Closing |
| -656.4 | 2.73113709 | BATF | Delayed Closing |
| -651.4 | 3.95853659 | OCT11 | Delayed Closing |
| -630.4 | 2.52126697 | AP-1 | Delayed Closing |
| -545.1 | 3.67901235 | OCT2 | Delayed Closing |
| -458.6 | 5.52631579 | CTCF | Delayed Closing |
| -456.6 | 4.01403509 | Bach2 | Delayed Closing |
| -376.4 | 3.82758621 | BORIS | Delayed Closing |
| -322.9 | 1.58471633 | Sox3 | Delayed Closing |
| -319 | 2.01482127 | Zic3 | Delayed Closing |
| -316.5 | 1.7875895 | TEAD1 | Delayed Closing |
| -312.3 | 1.84667115 | TEAD4 | Delayed Closing |
| -290.1 | 1.95287061 | TEAD | Delayed Closing |
| -277.3 | 1.66961959 | TEAD3 | Delayed Closing |
| -264.9 | 1.7847841 | Zic | Delayed Closing |
| -257.9 | 2.01147028 | TEAD2 | Delayed Closing |
| -1695 | 10.8218391 | OCT4-SOX2-TCF-NANOG | Early Transient |
| -911.5 | 2.64480472 | Zic3 | Early Transient |
| -742.9 | 1.9586743 | Sox3 | Early Transient |
| -716.2 | 3.64391144 | OCT4 | Early Transient |
| -710.7 | 2.4132948 | Unknown-ESC-element | Early Transient |
| -679.2 | 2.23247232 | Zic | Early Transient |
| -651.3 | 3.79431072 | OCT6 | Early Transient |
| -624.2 | 4.40123457 | Brn1 | Early Transient |
| -590 | 1.87987988 | Sox10 | Early Transient |
| -538.4 | 2.31458843 | Sox2 | Early Transient |
| -454.4 | 1.808491 | Sox6 | Early Transient |
| -451.5 | 3.70381232 | OCT11 | Early Transient |

| | | | |
|---|---|---|---|
| -389.6 | 2.07322835 | Sox4 | Early Transient |
| -366.9 | 2.22312373 | Sox17 | Early Transient |
| -348.8 | 3.39318885 | OCT2 | Early Transient |
| -316.1 | 1.80680437 | Sox15 | Early Transient |
| -225 | 2.83692308 | BORIS | Early Transient |
| -214.6 | 2.69774011 | Zfp281 | Early Transient |
| -197.8 | 1.70188133 | Sox9 | Early Transient |
| -192.7 | 1.52030217 | Tcf12 | Early Transient |
| -182.1 | 3.26455026 | CTCF | Early Transient |
| -174.3 | 2.456 | Foxa3 | Early Transient |
| -151.9 | 1.74374374 | Foxa2 | Early Transient |
| -150.5 | 1.31877729 | Ascl1 | Early Transient |
| -148.1 | 1.40143369 | Ap4 | Early Transient |
| -1246 | 9.49350649 | OCT4-SOX2-TCF-NANOG | Gradual Closing |
| -835.9 | 7.66 | CTCF | Gradual Closing |
| -738.6 | 5.10181818 | BORIS | Gradual Closing |
| -639.4 | 3.7826087 | OCT6 | Gradual Closing |
| -637.8 | 3.5502008 | OCT4 | Gradual Closing |
| -598.8 | 4.31309904 | Brn1 | Gradual Closing |
| -586.4 | 2.17296223 | TEAD1 | Gradual Closing |
| -497 | 1.9774044 | TEAD3 | Gradual Closing |
| -478.2 | 2.10192445 | TEAD4 | Gradual Closing |
| -461.3 | 1.78619756 | Sox3 | Gradual Closing |
| -443.2 | 2.2731569 | TEAD | Gradual Closing |
| -420.6 | 3.68488746 | OCT11 | Gradual Closing |
| -410.3 | 2.33953998 | TEAD2 | Gradual Closing |
| -349.1 | 1.79490085 | Sp5 | Gradual Closing |
| -336.2 | 3.4 | OCT2 | Gradual Closing |
| -316.1 | 1.81242079 | Fli1 | Gradual Closing |
| -309 | 1.56860465 | ERG | Gradual Closing |
| -294.5 | 1.8239726 | Etv2 | Gradual Closing |
| -280.2 | 2.00486855 | EWS:ERG-fusion | Gradual Closing |
| -278.3 | 1.61842105 | ETV1 | Gradual Closing |
| -276.9 | 1.62214829 | Sox10 | Gradual Closing |
| -274.8 | 1.74937186 | ETS1 | Gradual Closing |
| -266.6 | 2.02628812 | KLF3 | Gradual Closing |
| -263.7 | 2.11678832 | EWS:FLI1-fusion | Gradual Closing |
| -261.7 | 1.6971831 | ETV4 | Gradual Closing |
| -1269 | 2.33985765 | Sox3 | 4.5-day Transient |

| | | | |
|---|---|---|---|
| -1102 | 2.30503381 | Sox10 | 4.5-day Transient |
| -1080 | 3.10671378 | Sox2 | 4.5-day Transient |
| -877.9 | 2.21721802 | Sox6 | 4.5-day Transient |
| -844.8 | 2.82122905 | Sox4 | 4.5-day Transient |
| -794 | 2.42909664 | Sox15 | 4.5-day Transient |
| -762.1 | 3.0596745 | Sox17 | 4.5-day Transient |
| -570.8 | 2.39738562 | Otx2 | 4.5-day Transient |
| -488.3 | 2.27561608 | Lhx2 | 4.5-day Transient |
| -486.9 | 2.29081295 | Sox9 | 4.5-day Transient |
| -434.2 | 1.85359902 | Lhx3 | 4.5-day Transient |
| -412.9 | 2.10409146 | Lhx1 | 4.5-day Transient |
| -357.8 | 1.84932777 | LXH9 | 4.5-day Transient |
| -347.8 | 2.48915401 | Dlx3 | 4.5-day Transient |
| -306.4 | 1.50849744 | Nkx6.1 | 4.5-day Transient |
| -246.2 | 1.93366619 | Zic3 | 4.5-day Transient |
| -242.5 | 1.52179487 | Isl1 | 4.5-day Transient |
| -239.8 | 1.67870201 | Tcf12 | 4.5-day Transient |
| -230 | 1.57984791 | Ap4 | 4.5-day Transient |
| -230 | 1.67941454 | Tcf21 | 4.5-day Transient |
| -225.2 | 1.8777079 | Unknown-ESC-element | 4.5-day Transient |
| -219.9 | 1.76703553 | Zic | 4.5-day Transient |
| -188.7 | 1.76801058 | Myf5 | 4.5-day Transient |
| -185.5 | 1.71317365 | MyoD | 4.5-day Transient |
| -172.2 | 1.91316527 | LEF1 | 4.5-day Transient |
| -1439 | 1.89492326 | Sox3 | Late Opening |
| -1356 | 1.92666667 | Sox10 | Late Opening |
| -1102 | 2.30439684 | Sox2 | Late Opening |
| -905 | 1.74181478 | Sox6 | Late Opening |
| -796.8 | 2.15160703 | Sox9 | Late Opening |
| -788.6 | 1.89222842 | Sox15 | Late Opening |
| -783.7 | 2.25069252 | Sox17 | Late Opening |
| -775.4 | 2.11406619 | Sox4 | Late Opening |
| -761.4 | 1.95871782 | TEAD1 | Late Opening |
| -728.1 | 1.84589331 | TEAD3 | Late Opening |
| -678.1 | 1.9977591 | TEAD4 | Late Opening |
| -630 | 6.60169492 | Rfx2 | Late Opening |
| -594 | 7.1122449 | RFX | Late Opening |
| -582.8 | 2.0208605 | TEAD | Late Opening |
| -546.4 | 1.76744186 | Lhx2 | Late Opening |

| | | | |
|---|---|---|---|
| -481.2 | 1.58559499 | LXH9 | Late Opening |
| -473.3 | 4.68823529 | X-box | Late Opening |
| -431.6 | 3.25739645 | Rfx1 | Late Opening |
| -421.8 | 1.62823726 | Lhx1 | Late Opening |
| -401 | 1.98982188 | TEAD2 | Late Opening |
| -398.8 | 1.4632785 | Lhx3 | Late Opening |
| -366.2 | 1.31169094 | Nkx6.1 | Late Opening |
| -327.6 | 2.40142096 | Rfx5 | Late Opening |
| -301.3 | 1.37398594 | Isl1 | Late Opening |
| -259 | 3.76351351 | PAX6 | Late Opening |
| -748.5 | 2.09595202 | Sox3 | 2-day Transient |
| -668.1 | 2.68493151 | Zic3 | 2-day Transient |
| -642.8 | 2.70476911 | Sox2 | 2-day Transient |
| -631.6 | 2.4407497 | Zic | 2-day Transient |
| -612.8 | 2.02962085 | Sox10 | 2-day Transient |
| -576.8 | 2.51818182 | Unknown-ESC-element | 2-day Transient |
| -547.1 | 2.03432003 | Sox6 | 2-day Transient |
| -456.9 | 2.65259117 | Sox17 | 2-day Transient |
| -455.6 | 2.15005663 | Sox15 | 2-day Transient |
| -445.1 | 2.36457565 | Sox4 | 2-day Transient |
| -288.7 | 2.01215395 | Sox9 | 2-day Transient |
| -247.1 | 1.94718555 | Otx2 | 2-day Transient |
| -199.6 | 3.98429319 | OCT4-SOX2-TCF-NANOG | 2-day Transient |
| -140.5 | 1.53445851 | Tcf12 | 2-day Transient |
| -137.7 | 2.60773481 | Brn1 | 2-day Transient |
| -136.3 | 1.46016323 | Ap4 | 2-day Transient |
| -127.3 | 2.15017065 | OCT4 | 2-day Transient |
| -124.9 | 1.51735941 | Tcf21 | 2-day Transient |
| -124.7 | 1.14525678 | SCL | 2-day Transient |
| -123.6 | 1.81681682 | LEF1 | 2-day Transient |
| -106.4 | 1.44613321 | Lhx3 | 2-day Transient |
| -102.8 | 2.10331384 | OCT6 | 2-day Transient |
| -94.25 | 1.51461632 | MyoD | 2-day Transient |
| -91.34 | 1.56290439 | Lhx2 | 2-day Transient |
| -91.21 | 1.54371585 | Myf5 | 2-day Transient |
| -1413 | 2.17396866 | Sox3 | Gradual Opening |
| -1243 | 2.89943074 | Sox2 | Gradual Opening |
| -1148 | 2.11062246 | Sox10 | Gradual Opening |
| -905.3 | 2.01762425 | Sox6 | Gradual Opening |

| | | | |
|---|---|---|---|
| -884.6 | 2.56257822 | Sox4 | Gradual Opening |
| -824.8 | 2.2156587 | Sox15 | Gradual Opening |
| -759.2 | 2.68421053 | Sox17 | Gradual Opening |
| -580 | 2.19725864 | Sox9 | Gradual Opening |
| -526.8 | 2.0541555 | Lhx2 | Gradual Opening |
| -480.6 | 1.72080166 | Lhx3 | Gradual Opening |
| -472.3 | 1.94969819 | Lhx1 | Gradual Opening |
| -420.6 | 1.74890916 | LXH9 | Gradual Opening |
| -342.9 | 2.1630149 | Dlx3 | Gradual Opening |
| -342.2 | 1.43294839 | Nkx6.1 | Gradual Opening |
| -331 | 1.88116459 | Otx2 | Gradual Opening |
| -298.9 | 5 | Rfx2 | Gradual Opening |
| -289.3 | 5.32786885 | RFX | Gradual Opening |
| -269.8 | 4.58823529 | PAX6 | Gradual Opening |
| -263 | 1.46126761 | Isl1 | Gradual Opening |
| -196.8 | 3.68539326 | X-box | Gradual Opening |
| -144.9 | 1.40449853 | Ap4 | Gradual Opening |
| -140.2 | 2.38258575 | Rfx1 | Gradual Opening |
| -130.1 | 1.65622424 | LEF1 | Gradual Opening |
| -99.23 | 1.44748858 | MyoD | Gradual Opening |
| -99.14 | 1.38705234 | Tcf21 | Gradual Opening |

**Table S4:** Full list of TF displaying differential expression during the time course clustered by expression pattern, as visualized in Figure 4C**.**

| Transcription Factor | Cluster number |
|---|---|
| ZNF684 | 1 |
| MIER1 | 1 |
| LHX8 | 1 |
| NHLH2 | 1 |
| NR5A2 | 1 |
| TFB2M | 1 |
| ZNF124 | 1 |
| CEBPZ | 1 |
| BOLA3 | 1 |
| KCMF1 | 1 |
| LRRFIP1 | 1 |
| ZNF860 | 1 |
| ZNF35 | 1 |
| HLTF | 1 |
| SKIL | 1 |
| ZNF876P | 1 |
| LYAR | 1 |
| NFXL1 | 1 |

| | |
|---|---|
| ZFP42 | 1 |
| NR3C1 | 1 |
| ZNF165 | 1 |
| POU5F1 | 1 |
| ZBTB24 | 1 |
| HEY2 | 1 |
| NCOA7 | 1 |
| ETV1 | 1 |
| NFE2L3 | 1 |
| DNAJC2 | 1 |
| ZNF398 | 1 |
| ZMAT4 | 1 |
| THAP1 | 1 |
| TCF24 | 1 |
| PRDM14 | 1 |
| MSC | 1 |
| TERF1 | 1 |
| HNF4G | 1 |
| RUNX1T1 | 1 |
| POU5F1B | 1 |
| ZNF483 | 1 |
| MKX | 1 |
| ZNF239 | 1 |
| ZNF485 | 1 |
| TFAM | 1 |
| NOC3L | 1 |
| PCGF6 | 1 |
| VENTX | 1 |
| ZNF195 | 1 |
| ESRRA | 1 |
| ETS1 | 1 |
| ZBTB44 | 1 |
| NANOG | 1 |
| BHLHE41 | 1 |
| ARNTL2 | 1 |
| ATF1 | 1 |
| YEATS4 | 1 |
| PAWR | 1 |
| GTF3A | 1 |
| FOXN3 | 1 |
| ZNF770 | 1 |
| ONECUT1 | 1 |
| ZNF267 | 1 |
| SNAI3 | 1 |
| ZNF232 | 1 |
| PHB | 1 |
| SMAD7 | 1 |
| ZNF57 | 1 |
| ZNF121 | 1 |

| | |
|---|---|
| ZNF878 | 1 |
| ZNF90 | 1 |
| ZNF146 | 1 |
| ZNF114 | 1 |
| ZNF649 | 1 |
| ZNF836 | 1 |
| ZNF28 | 1 |
| ZNF468 | 1 |
| ZNF845 | 1 |
| ZNF761 | 1 |
| ZNF813 | 1 |
| TCF15 | 1 |
| OVOL2 | 1 |
| PHF5A | 1 |
| ZNF670 | 2 |
| MYT1L | 2 |
| ATF2 | 2 |
| ZNF385D | 2 |
| ETV5 | 2 |
| LCORL | 2 |
| RBPJ | 2 |
| CLOCK | 2 |
| ZNF354A | 2 |
| ZNF184 | 2 |
| RUNX2 | 2 |
| ZUFSP | 2 |
| SNAI2 | 2 |
| RORB | 2 |
| ZBTB6 | 2 |
| ZNF143 | 2 |
| THAP2 | 2 |
| ZNF605 | 2 |
| ZFP1 | 2 |
| IRF8 | 2 |
| ZNF624 | 2 |
| KAT7 | 2 |
| ZNF555 | 2 |
| ZNF566 | 2 |
| ZNF225 | 2 |
| ZNF234 | 2 |
| ZNF615 | 2 |
| ZNF415 | 2 |
| ZNF765 | 2 |
| ID1 | 2 |
| TP73 | 3 |
| ENO1 | 3 |
| ID3 | 3 |
| GRHL3 | 3 |
| ZNF593 | 3 |

| | |
|---|---|
| ZC3H12A | 3 |
| FOXE3 | 3 |
| FOXD3 | 3 |
| TTF2 | 3 |
| ZNF697 | 3 |
| ZBTB7B | 3 |
| ZNF281 | 3 |
| ELF3 | 3 |
| IRF6 | 3 |
| ATF3 | 3 |
| BATF3 | 3 |
| MYCN | 3 |
| PREB | 3 |
| EPAS1 | 3 |
| BCL11A | 3 |
| EMX1 | 3 |
| TCF7L1 | 3 |
| FOXI3 | 3 |
| TFCP2L1 | 3 |
| NFE2L2 | 3 |
| SP110 | 3 |
| SP140 | 3 |
| GBX2 | 3 |
| HES6 | 3 |
| THAP4 | 3 |
| BHLHE40 | 3 |
| CSRNP1 | 3 |
| NFKBIZ | 3 |
| FOXL2 | 3 |
| AHRR | 3 |
| IRX4 | 3 |
| ZNF622 | 3 |
| IRF1 | 3 |
| MXD3 | 3 |
| IRF4 | 3 |
| JARID2 | 3 |
| ZFP57 | 3 |
| TCF19 | 3 |
| HMGA1 | 3 |
| TFEB | 3 |
| NFKBIE | 3 |
| ZBTB2 | 3 |
| MAFK | 3 |
| AHR | 3 |
| PARP12 | 3 |
| NKX3-1 | 3 |
| CEBPD | 3 |
| GRHL2 | 3 |
| TRPS1 | 3 |

| | |
|---|---|
| ZHX2 | 3 |
| MYC | 3 |
| MAFA | 3 |
| SMARCA2 | 3 |
| NFIB | 3 |
| BNC2 | 3 |
| NFX1 | 3 |
| PAX5 | 3 |
| LHX6 | 3 |
| ZNF488 | 3 |
| NFKB2 | 3 |
| NKX1-2 | 3 |
| FOXI2 | 3 |
| UTF1 | 3 |
| ARNTL | 3 |
| E2F8 | 3 |
| ZBTB3 | 3 |
| OVOL1 | 3 |
| FOSL1 | 3 |
| PGR | 3 |
| ZNF202 | 3 |
| TEAD4 | 3 |
| PHB2 | 3 |
| YBX3 | 3 |
| VDR | 3 |
| POU6F1 | 3 |
| DDIT3 | 3 |
| NOC4L | 3 |
| PSPC1 | 3 |
| FOXO1 | 3 |
| TFDP1 | 3 |
| JDP2 | 3 |
| ZBTB42 | 3 |
| BNC1 | 3 |
| ZSCAN2 | 3 |
| MESP2 | 3 |
| ZSCAN10 | 3 |
| ZNF469 | 3 |
| TAX1BP3 | 3 |
| YBX2 | 3 |
| SOX15 | 3 |
| MLX | 3 |
| ETV4 | 3 |
| UBTF | 3 |
| SP6 | 3 |
| NFE2L1 | 3 |
| SMARCD2 | 3 |
| FOXJ1 | 3 |
| MAFG | 3 |

| | |
|---|---|
| TGIF1 | 3 |
| KLF16 | 3 |
| NR2F6 | 3 |
| ZNF101 | 3 |
| TSHZ3 | 3 |
| DPF1 | 3 |
| NFKBIB | 3 |
| ZNF296 | 3 |
| FOXA3 | 3 |
| IRF3 | 3 |
| ATF5 | 3 |
| SPIB | 3 |
| ZNF134 | 3 |
| ZNF552 | 3 |
| FOXA2 | 3 |
| DNMT3B | 3 |
| E2F1 | 3 |
| MYBL2 | 3 |
| CEBPB | 3 |
| ZFP64 | 3 |
| OLIG2 | 3 |
| OLIG1 | 3 |
| ERG | 3 |
| XBP1 | 3 |
| MAFF | 3 |
| ATF4 | 3 |
| TFE3 | 3 |
| POU3F4 | 3 |
| ELF4 | 3 |
| MYCL | 4 |
| DMBX1 | 4 |
| ETV3L | 4 |
| PBX1 | 4 |
| RXRG | 4 |
| PRRX1 | 4 |
| MTA3 | 4 |
| SP5 | 4 |
| THRB | 4 |
| ZBTB47 | 4 |
| HESX1 | 4 |
| MYB | 4 |
| OLIG3 | 4 |
| CREB3L2 | 4 |
| ASH2L | 4 |
| TOX | 4 |
| FOXH1 | 4 |
| FOXB2 | 4 |
| LMX1B | 4 |
| BARHL1 | 4 |

| | |
|---|---|
| HHEX | 4 |
| PAX2 | 4 |
| HMX3 | 4 |
| HMX2 | 4 |
| CREB3L1 | 4 |
| BATF2 | 4 |
| PHOX2A | 4 |
| POU2F3 | 4 |
| PKNOX2 | 4 |
| RARG | 4 |
| NFE2 | 4 |
| STAT6 | 4 |
| GLI1 | 4 |
| ZIC5 | 4 |
| ZIC2 | 4 |
| NKX2-8 | 4 |
| DPF3 | 4 |
| CEBPA | 4 |
| CRX | 4 |
| SNAI1 | 4 |
| TFAP2C | 4 |
| CTCFL | 4 |
| PCBP3 | 4 |
| FOXO4 | 4 |
| MBNL3 | 4 |
| ZIC3 | 4 |
| HES3 | 5 |
| ZBTB17 | 5 |
| ZNF436 | 5 |
| E2F2 | 5 |
| ZNF362 | 5 |
| ZSCAN20 | 5 |
| POU3F1 | 5 |
| FOXO6 | 5 |
| ZCCHC11 | 5 |
| ALX3 | 5 |
| PHTF1 | 5 |
| GABPB2 | 5 |
| RFX5 | 5 |
| GATAD2B | 5 |
| ASH1L | 5 |
| IFI16 | 5 |
| LMX1A | 5 |
| TBX19 | 5 |
| ZBTB41 | 5 |
| LHX9 | 5 |
| ZC3H11A | 5 |
| ELK4 | 5 |
| RCOR3 | 5 |

| | |
|---|---|
| SOX11 | 5 |
| ID2 | 5 |
| ZNF512 | 5 |
| SIX3 | 5 |
| REL | 5 |
| OTX1 | 5 |
| ZNF514 | 5 |
| ZC3H8 | 5 |
| ZC3H6 | 5 |
| ZEB2 | 5 |
| MBD5 | 5 |
| CSRNP3 | 5 |
| SP9 | 5 |
| ZNF385B | 5 |
| ZNF804A | 5 |
| SATB1 | 5 |
| NR1D2 | 5 |
| ZNF445 | 5 |
| PCBP4 | 5 |
| ZNF148 | 5 |
| TFDP2 | 5 |
| MECOM | 5 |
| SOX2 | 5 |
| BCL6 | 5 |
| HES1 | 5 |
| MXD4 | 5 |
| ZNF518B | 5 |
| HOPX | 5 |
| REST | 5 |
| NKX6-1 | 5 |
| TET2 | 5 |
| NEUROG2 | 5 |
| IRX1 | 5 |
| ZNF608 | 5 |
| ZFP2 | 5 |
| ZNF879 | 5 |
| FOXC1 | 5 |
| SOX4 | 5 |
| ZSCAN9 | 5 |
| ZSCAN26 | 5 |
| ZKSCAN3 | 5 |
| ZSCAN23 | 5 |
| PBX2 | 5 |
| ZBTB22 | 5 |
| ZNF451 | 5 |
| ZNF292 | 5 |
| HSF2 | 5 |
| MLLT4 | 5 |
| ZNF853 | 5 |

| | |
|---|---|
| POU6F2 | 5 |
| ZNF713 | 5 |
| ZNF117 | 5 |
| ZNF789 | 5 |
| ZKSCAN1 | 5 |
| CUX1 | 5 |
| KMT2E | 5 |
| HBP1 | 5 |
| FEZF1 | 5 |
| ZNF467 | 5 |
| SMARCD3 | 5 |
| KMT2C | 5 |
| ZNF395 | 5 |
| HMBOX1 | 5 |
| PURG | 5 |
| PLAG1 | 5 |
| ZFHX4 | 5 |
| ZBTB10 | 5 |
| GLI4 | 5 |
| ZNF517 | 5 |
| ZNF250 | 5 |
| RFX3 | 5 |
| MLLT3 | 5 |
| ZFP37 | 5 |
| ZBTB26 | 5 |
| NR5A1 | 5 |
| NR6A1 | 5 |
| PBX3 | 5 |
| PRRX2 | 5 |
| ZNF438 | 5 |
| ZNF248 | 5 |
| ZNF25 | 5 |
| MXI1 | 5 |
| ASCL2 | 5 |
| ZNF214 | 5 |
| TUB | 5 |
| DBX1 | 5 |
| KMT2A | 5 |
| SOX5 | 5 |
| ZNF641 | 5 |
| ZNF740 | 5 |
| SP7 | 5 |
| ZC3H10 | 5 |
| SMARCC2 | 5 |
| STAT2 | 5 |
| NR2C1 | 5 |
| LHX5 | 5 |
| ZNF664 | 5 |
| ZNF84 | 5 |

| | |
|---|---|
| TSC22D1 | 5 |
| DACH1 | 5 |
| SOX21 | 5 |
| ZNF219 | 5 |
| HOMEZ | 5 |
| NFATC4 | 5 |
| FOXA1 | 5 |
| OTX2 | 5 |
| ARID4A | 5 |
| SIX6 | 5 |
| VSX2 | 5 |
| FOS | 5 |
| ESRRB | 5 |
| GSC | 5 |
| ZNF280D | 5 |
| FOXB1 | 5 |
| SMAD6 | 5 |
| SCAPER | 5 |
| ZNF710 | 5 |
| ZNF785 | 5 |
| TOX3 | 5 |
| IRX3 | 5 |
| IRX5 | 5 |
| ZFP90 | 5 |
| ZNF19 | 5 |
| MAF | 5 |
| ZBTB4 | 5 |
| ZNF287 | 5 |
| MLLT6 | 5 |
| THRA | 5 |
| ZNF385C | 5 |
| STAT5B | 5 |
| EZH1 | 5 |
| DLX3 | 5 |
| VEZF1 | 5 |
| ZNF519 | 5 |
| ZNF521 | 5 |
| SETBP1 | 5 |
| SMAD2 | 5 |
| ZBTB7C | 5 |
| TCF4 | 5 |
| RAX | 5 |
| ZNF516 | 5 |
| SALL3 | 5 |
| ZNF558 | 5 |
| ZNF699 | 5 |
| ZNF559-ZNF177 | 5 |
| ZNF177 | 5 |
| ZNF266 | 5 |

| | |
|---|---|
| ZNF846 | 5 |
| ZNF653 | 5 |
| ZNF441 | 5 |
| ZNF491 | 5 |
| ZNF440 | 5 |
| ZNF439 | 5 |
| ZNF69 | 5 |
| ZNF763 | 5 |
| ZNF433 | 5 |
| ZNF709 | 5 |
| JUND | 5 |
| PBX4 | 5 |
| ZNF737 | 5 |
| ZNF91 | 5 |
| ZFP14 | 5 |
| ZNF529 | 5 |
| ZNF568 | 5 |
| ZNF585A | 5 |
| HKR1 | 5 |
| ZNF793 | 5 |
| ZNF571 | 5 |
| ZFP30 | 5 |
| ZNF781 | 5 |
| ZNF546 | 5 |
| ZNF780B | 5 |
| ZNF428 | 5 |
| ZNF155 | 5 |
| ZNF224 | 5 |
| ZNF226 | 5 |
| ZNF112 | 5 |
| ZNF285 | 5 |
| SIX5 | 5 |
| ZNF432 | 5 |
| ZNF610 | 5 |
| ZNF880 | 5 |
| ZNF83 | 5 |
| ZNF611 | 5 |
| ZNF702P | 5 |
| ZNF579 | 5 |
| ZNF580 | 5 |
| ZNF419 | 5 |
| ZNF211 | 5 |
| ZNF606 | 5 |
| ZNF135 | 5 |
| ZSCAN18 | 5 |
| ZNF329 | 5 |
| ZNF446 | 5 |
| MZF1 | 5 |
| ZNF343 | 5 |

| | |
|---|---|
| C20orf194 | 5 |
| BMP2 | 5 |
| TGIF2 | 5 |
| ZHX3 | 5 |
| ZNF334 | 5 |
| NCOA3 | 5 |
| SALL4 | 5 |
| MYT1 | 5 |
| ZNF280A | 5 |
| ZNF70 | 5 |
| SOX10 | 5 |
| TEF | 5 |
| KLF8 | 5 |
| ATRX | 5 |
| ZNF711 | 5 |
| SMARCA1 | 5 |
| ZNF449 | 5 |
| HES4 | 6 |
| HES5 | 6 |
| PRDM16 | 6 |
| CASZ1 | 6 |
| HEYL | 6 |
| HIVEP3 | 6 |
| DMRTA2 | 6 |
| JUN | 6 |
| NHLH1 | 6 |
| MAEL | 6 |
| ZBTB37 | 6 |
| PROX1 | 6 |
| ESRRG | 6 |
| ZBTB18 | 6 |
| FOSL2 | 6 |
| MEIS1 | 6 |
| MXD1 | 6 |
| VAX2 | 6 |
| ATOH8 | 6 |
| AFF3 | 6 |
| POU3F3 | 6 |
| IKZF2 | 6 |
| EOMES | 6 |
| FEZF2 | 6 |
| ATXN7 | 6 |
| MITF | 6 |
| ZNF654 | 6 |
| ZIC4 | 6 |
| ZIC1 | 6 |
| MSX1 | 6 |
| NKX3-2 | 6 |
| KLF3 | 6 |

| | |
|---|---|
| NFKB1 | 6 |
| LEF1 | 6 |
| MEF2C | 6 |
| NR2F1 | 6 |
| EGR1 | 6 |
| EBF1 | 6 |
| TFAP2A | 6 |
| ID4 | 6 |
| ATF6B | 6 |
| RXRB | 6 |
| TEAD3 | 6 |
| BACH2 | 6 |
| POU3F2 | 6 |
| SIM1 | 6 |
| HIVEP2 | 6 |
| PLAGL1 | 6 |
| UNCX | 6 |
| FOXK1 | 6 |
| ZNF316 | 6 |
| SP8 | 6 |
| CREB5 | 6 |
| GLI3 | 6 |
| DLX5 | 6 |
| FOXP2 | 6 |
| ZNF800 | 6 |
| EGR3 | 6 |
| EBF2 | 6 |
| ZNF703 | 6 |
| KAT6A | 6 |
| ST18 | 6 |
| BHLHE22 | 6 |
| HEY1 | 6 |
| ZNF704 | 6 |
| ZNF706 | 6 |
| ZHX1 | 6 |
| SCRT1 | 6 |
| GLIS3 | 6 |
| DMRTA1 | 6 |
| ZBTB5 | 6 |
| NR4A3 | 6 |
| LHX2 | 6 |
| ZEB1 | 6 |
| ARID5B | 6 |
| ZNF503 | 6 |
| VAX1 | 6 |
| EMX2 | 6 |
| EBF3 | 6 |
| SOX6 | 6 |
| PAX6 | 6 |

| | |
|---|---|
| ZBTB16 | 6 |
| NR4A1 | 6 |
| ZNF385A | 6 |
| NEUROD4 | 6 |
| RFX4 | 6 |
| FOXN4 | 6 |
| SMAD9 | 6 |
| SOX1 | 6 |
| FOXG1 | 6 |
| NPAS3 | 6 |
| ELMSAN1 | 6 |
| MEIS2 | 6 |
| SMAD3 | 6 |
| SKOR1 | 6 |
| ARNT2 | 6 |
| ZNF774 | 6 |
| NR2F2 | 6 |
| ZNF843 | 6 |
| ZFHX3 | 6 |
| FOXC2 | 6 |
| FOXL1 | 6 |
| CAMTA2 | 6 |
| HES7 | 6 |
| SOX9 | 6 |
| ONECUT2 | 6 |
| TSHZ1 | 6 |
| RFX2 | 6 |
| ZNF333 | 6 |
| ZNF536 | 6 |
| ZNF599 | 6 |
| ZNF382 | 6 |
| FOSB | 6 |
| MEIS3 | 6 |
| ZNF808 | 6 |
| ZNF264 | 6 |
| ZSCAN1 | 6 |
| ZNF584 | 6 |
| ZNF132 | 6 |
| EBF4 | 6 |
| INSM1 | 6 |
| ZNF337 | 6 |
| MAFB | 6 |
| TSHZ2 | 6 |
| SIM2 | 6 |
| PPARA | 6 |
| ARX | 6 |
| DACH2 | 6 |
| ZMAT1 | 6 |
| BHLHB9 | 6 |

| | |
|---|---|
| ZNF75D | 6 |
| SOX3 | 6 |
| MECP2 | 6 |
